# The NAC38-NAC58-FUSCA3 module regulates outer integument differentiation in the seed coat of *Arabidopsis thaliana*

**DOI:** 10.64898/2026.11.23.762040

**Authors:** Lin Wu, Jasmin Patel, Yang Yang, Yunling Ye, Dianqiu Lv, Jian Wu, Richard Yang, Maria Barsamyan, Myles Matundan, Marc Mohammadi, Isabel Molina, Sonia Gazzarrini

**Author notes:** These authors contributed equally to this paper (Lin Wu, Jasmin Patel).

## Abstract

The seed coat protects the embryo from environmental stresses while controlling important agricultural traits, such as seed dormancy, germination, and longevity. The Arabidopsis seed coat epidermal layer secretes a pectin-rich cell wall, the mucilage, which regulates seed hydration and germination. In the subepidermal layer, the lipid polyester suberin is deposited in the cell wall, where it forms a protective barrier against solutes and gases. Currently, the mechanisms coordinating the development of these important cell layers of the outer integument are poorly understood. Here, we show that *NAC38* and *NAC58* are preferentially expressed in the outer integument and directly promote the expression of *FUSCA3 (FUS3)*, a master regulator of seed maturation that promotes fatty acid biosynthesis and desiccation tolerance. In a feedback loop, FUS3 positively regulates *NAC38* and *NAC58*. Loss-of-function mutations in *NAC38, NAC58,* and *FUS3* transcription factors affect the development of the outer integument, by reducing radial wall thickening and mucilage extrusion, while altering suberin monomer levels. Loss of *FUS3* also decreases total suberin content, suggesting a prominent role for FUS3 in this process. During prolonged dry storage, loss of *NAC38* decreases seedling growth and loss of *NAC38* and *NAC58* increase seed sensitivity to salinity stress. Genome-wide binding sites and transcriptomic analysis support a role for these transcription factors in cell wall-related processes, by directly regulating mucilage- and suberin-related genes, through shared and distinct gene sets. This study uncovered novel regulators of mucilage secretory cell differentiation and suberin monomers accumulation, which affect seed longevity and salt sensitivity of aged seeds.

## INTRODUCTION

In flowering plants, double fertilization initiates seed development, giving rise to three tissues: the embryo, endosperm, and seed coat (SC). The maternally derived SC (testa) protects the developing embryo against mechanical, biotic, and abiotic stresses, while also influencing seed size, longevity, dispersal, dormancy, and germination (Haughn and Chaudhury, 2005; Radchuk and Borisjuk, 2014; Francoz et al., 2015; Ezquer et al., 2016; Figueiredo et al., 2016). The structural and chemical properties of the SC are critical for embryo survival under unfavourable conditions, underpinning agricultural efficiency and food security.

In *Arabidopsis thaliana*, the SC develops from the outer and inner integuments of the ovule, which differentiate into specialized cell layers after fertilization (Beeckman et al., 2000; Windsor et al., 2000; Haughn and Chaudhury, 2005). The outermost integument layer (oi2) forms the seed coat epidermis, which secretes mucilage—a polysaccharide-rich specialized cell wall. Upon hydration, mucilage expands and extrudes by rupturing the primary cell wall, promoting germination, preventing desiccation, and aiding seed dispersal (Penfield et al., 2001; Debeaujon et al., 2003; Rautengarten et al., 2008; Arsovski et al., 2010; Harpaz-Saad et al., 2011; Western, 2012; Yang et al., 2012; Francoz et al., 2015). A three-tier MYB-bHLH-WD40 (MBW) cascade regulates SC development. The tier 3 MBW complex and APETALA2 (AP2) promote mucilage secretory cell differentiation and mucilage production by positively regulating tier-2 TFs (Golz et al., 2018; Huang et al., 2023; Xu et al., 2023). Tier-3 and tier-2 TFs also act with tier-1 TFs to control the expression of structural genes (Baudry et al., 2004; Lepiniec et al., 2006; Li et al., 2009; Arsovski et al., 2010; Khan et al., 2014; Francoz et al., 2015; Li et al., 2020).

The subepidermal layer (oi1) undergoes cell wall thickening and deposits lignin and suberin. Lignin provides mechanical resistance and influences seed permeability, germination, and cold stress response (Vishwanath et al., 2013; Francoz et al., 2018; Menino et al., 2023; Hyvärinen et al., 2025; Zhuang et al., 2025). Suberin forms lamellae in the oi1 and chalaza SC, limiting permeability and promoting dormancy (Beisson et al., 2007; Molina et al., 2009; Vishwanath et al., 2013; Yadav et al., 2014; Fedi et al., 2017; Gou et al., 2017; de Silva et al., 2021; Kosma et al., 2025). MYB9 and MYB107 positively regulate seed coat suberin biosynthesis (Roscoe et al., 2015; Lashbrooke et al., 2016; Kosma et al., 2025). While suberin patterning is well studied in roots (Serra and Geldner, 2022), mechanisms regulating suberin deposition in seeds remain largely unknown and defining spatiotemporal functions of these regulators in adjacent SC layers remains challenging.

The B3-domain transcription factor (TF) *FUSCA3* (*FUS3*) is a master regulator of seed maturation. FUS3 promotes storage proteins and lipids (triacylglycerol) accumulation, and establishment of dormancy and desiccation tolerance, partly via opposing regulation of ABA and GA pathways (Keith et al., 1994; Meinke et al., 1994; Curaba et al., 2004; Gazzarrini et al., 2004; Tiedemann et al., 2008; Roscoe et al., 2015). Seeds of the *fus3* loss-of-function mutant are desiccation intolerant and unviable at maturity and show approximately one-third of wild-type total fatty acids (Keith et al., 1994; (Meinke et al., 1994; Tiedemann et al., 2008; Roscoe et al., 2015). In agreement with this, transcriptomic data show that FUS3 controls primary metabolism genes, including *FAE1/KCS18*, essential for the synthesis of very long-chain fatty acids (C20–C22), which are precursors of TAGs but also wax and suberin (Yamamoto et al., 2010; Wang and Perry, 2013; Batsale et al., 2021). Post-embryonic and ectopic *FUS3* expression activates TAG pathway genes and increases TAG content, underscoring its prominent role in this pathway (Zhang et al., 2016).

While FUS3 function is well studied in embryos, *FUS3* is also expressed in the seed coat (SC), where its role remains largely unknown despite FUS3 role in desiccation tolerance and dormancy (Wu et al., 2020). Here, we identified two NAC domain TFs, NAC38 and NAC58, that are preferentially expressed in the outer integument of the SC and directly bind to and induce *FUS3* expression. The large family of NAC TFs (more than 100 in Arabidopsis) are important regulators of secondary cell wall biosynthesis and stress responses (Singh et al., 2021; Han et al., 2023; Dong et al., 2024). Loss-of-function mutants in *NAC* and *FUS3* affect mucilage secretory cell development, including mucilage structure and extrusion, and suberin monomer accumulation, with FUS3 playing a more prominent role in total suberin accumulation in seeds. While germination and viability of *fus3* loss-of-function mutant seeds dramatically decrease soon after dry storage, loss of *NAC38* and *NAC58* delay seedling establishment in seeds after prolonged dry storage, with loss of *NAC38* causing a decrease in seed longevity. Furthermore, loss of *NAC38* and *NAC58* increase seed sensitivity to salinity stress after prolonged dry storage. Genome-wide identification of target genes directly controlled by these TFs shows that NAC38, NAC58 and FUS3 directly and indirectly regulate secondary metabolism and cell wall-related processes, including mucilage and suberin pathways, through shared as well as distinct gene sets. This study uncovers a novel transcriptional module important for proper development of the outer SC integument in Arabidopsis, which have implications in seed longevity and sensitivity to salinity stress.

## RESULTS

### NAC38 and NAC58 directly associate with the *FUS3* genomic locus and positively regulated *FUS3* expression

In the seed coat (SC), *FUS3* expression starts right after fertilization, but its function and regulation in this tissue, which is important for dormancy and embryo protection, is unknown (Wu et al., 2020). To identify transcriptional regulators of *FUS3*, a 500 bp fragment from -1000 to -501 bp of *FUS3* promoter was used in Y1H as the bait against an Arabidopsis transcription factor (TF) library (∼1400 TFs) (Mitsuda et al., 2010). Sequencing results indicated that the NAC58 transcription factor could directly bind to the *FUS3* promoter (**Figure 1A**). Accordingly, several *cis* elements previously shown to be bound by different NACs are found in the *FUS3* promoter from -500 to -1000 (Tran et al., 2004; Olsen et al., 2005; Xu et al., 2013; Lindemose et al., 2014) (**Supplementary Figure 1A**). Phylogenetic analysis of the NAC transcription factors in Arabidopsis showed that NAC38 is the closest relative of NAC58 (Takahashi et al., 2019) and their N-terminal NAC domains show high amino acid sequence conservation, while the C-terminus is more divergent (**Supplementary Figure 1C and D**). Although no binding was observed between NAC38 and the *FUS3* promoter (-1000 to -501) in Y1H (**Figure 1A**), analysis of DNA-affinity purification sequencing (DAP-seq) (O’Malley et al., 2016) revealed that both NAC38 and NAC58 could bind to the DNA fragment between the fifth exon, the fifth intron, and the sixth exon of the *FUS3* genomic locus (**Figure 1B**). Consistently, sequence analysis showed three core NAC motifs (5’-CACG-3’) (Tran et al., 2004) in this fragment (**Supplementary Figure 1B**). Y1H confirmed binding of NAC38/58 to the *FUS3* genomic locus (1125 to 1851 bp; **Figure 1C**).

**Figure 1.**
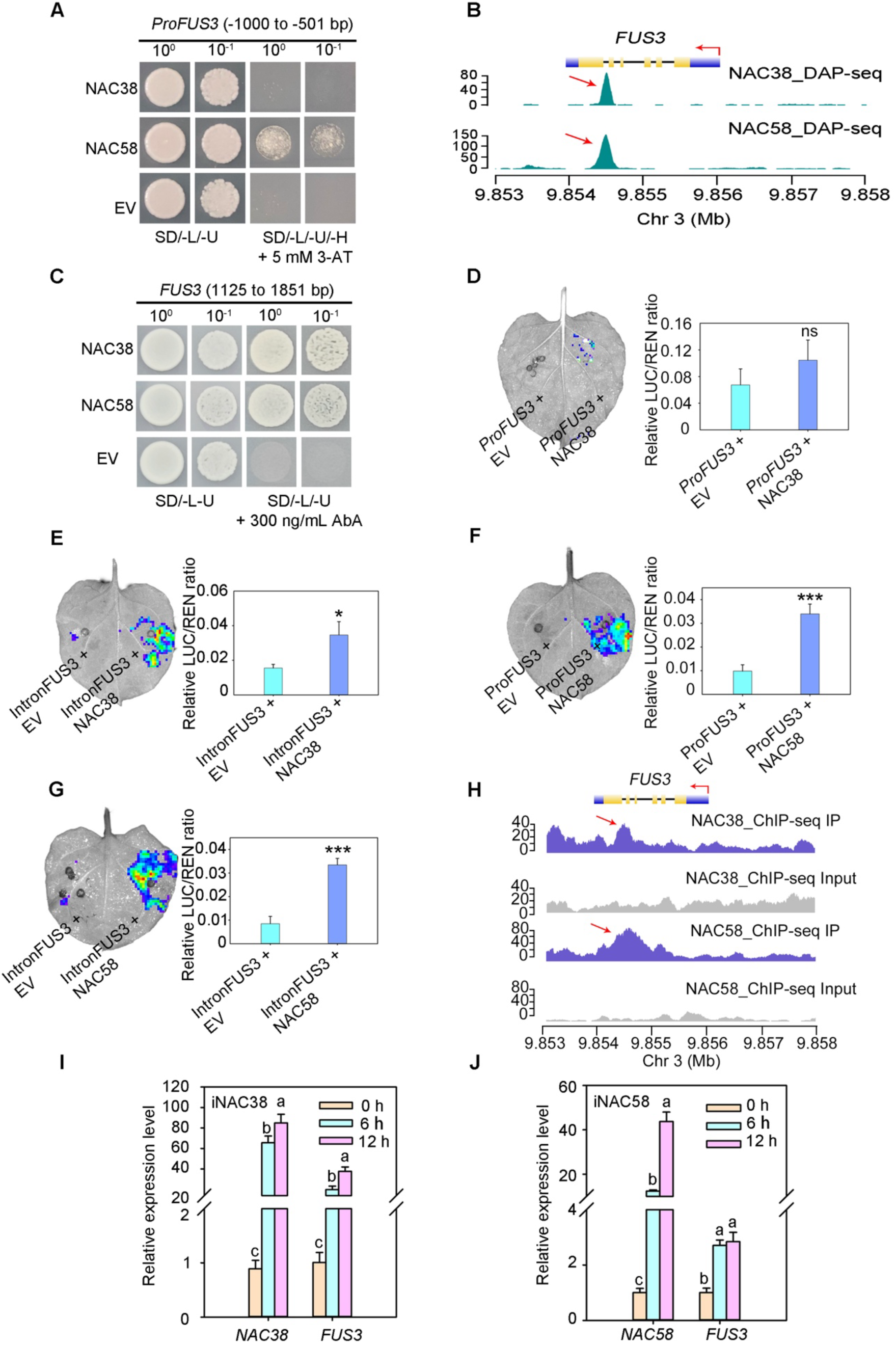
NAC38 and NAC58 bind to the promoter and genomic locus of *FUS3* and induce its expression. **A**, NAC58 binds to the *FUS3* promoter region (–1000 to –501 bp) in yeast one-hybrid (Y1H) assays. Colonies were selected on synthetic dropout medium lacking uracil, leucine, and histidine (SD–Ura–Leu–His) supplemented with 5 mM 3-amino-1,2,4-triazole (3-AT). “Ev” denotes the empty vector control; “10^-1^” indicates a 10-fold dilution of the initial culture (OD_600_ = 1.0). **B**, Both NAC38 and NAC58 directly bind to the *FUS3* genomic locus (1125–1851 bp), as determined by DNA affinity purification sequencing (DAP-seq). DAP-seq data are from O’Malley et al. (2016); visualization was performed using the AnnoJ browser of the Plant Cistrome Database (http://neomorph.salk.edu/dap_web/pages/index.php). The blue arrow represents the transcription direction of *FUS3*. The red arrows represent the regions bound by NAC38 or NAC58. **C**, NAC38 and NAC58 also bind to the *FUS3* genomic locus (1125–1851 bp) in yeast one-hybrid (H1Y) assays. Colonies were selected on SD–Ura–Leu medium supplemented with 300 ng/mL aureobasidin A (AbA). **D–G**, Both NAC38 and NAC58 function as transcriptional activators of *FUS3*. In dual-luciferase reporter assays conducted in *Nicotiana benthamiana* leaves, either the *FUS3* promoter fragment (–1000 to –501 bp) or the genomic locus (1125–1851 bp) was co-infiltrated with either an empty effector vector (EV) or effector constructs expressing NAC38 (**D, E**) or NAC58 (**F, G**). Luminescence was measured three days post-infiltration, and results are presented as the relative LUC/REN ratio. Data represent means ± SD from three independent biological replicates. Student’s *t*-test (* *P* < 0.05; *** *P* < 0.001); ns, no statistically significant difference. **H**, Chromatin immunoprecipitation sequencing (ChIP-seq) reveals *in vivo* occupancy of NAC38 and NAC58 at the *FUS3* locus, visualized using the Integrative Genomics Viewer (IGV). The blue arrow represents the transcription direction of *FUS3*. The red arrows represent the regions bound by NAC38 or NAC58. **I, J**, Induction of *NAC38* (**I)** or *NAC58* (**J**) expression in respective β-estradiol-inducible lines (*iNAC38* and *iNAC58*) leads to significant upregulation of *FUS3* transcript levels, as confirmed by quantitative RT-PCR. Different letters indicate statistically significant differences determined by Student’s *t*-test (*P* < 0.05).

To confirm these results *in planta*, a transient dual-LUC reporter assay was performed in *N. benthamiana* leaves. A significant enhancement in the luciferase activities driven by the *FUS3* promoter (-1000 to -501 bp) or *FUS3* genomic fragment (1125 to 1851 bp) when NAC38/58 were transiently expressed compared to the control (EV), with the exception of the combination of NAC38 and *FUS3* promoter (**Figure 1D to 1G**). To determine whether NAC38/58 also bind to the *FUS3* locus *in vivo*, we generated inducible overexpression lines, *iNAC38-HA-flag* (*iNAC38*) and *iNAC58-HA-flag* (*iNAC58*), under the control of a β–estradiol inducible promoter. Chromatin immunoprecipitation sequencing (ChIP-seq) showed that both NACs directly bind to the *FUS3* locus *in vivo* (**Figure 1H**). RT-qPCR results confirmed that transcripts of *FUS3* were strongly induced by iNAC38 and slightly upregulated by iNAC58 after a short treatment with β–estradiol (**Figure 1I and 1J**).

Taken together, these results indicated that NAC38 and NAC58 directly bind to the *FUS3* promoter or the genomic locus and positively regulate *FUS3* expression.

### *FUS3, NAC38*, and *NAC58* are co-expressed in the seed coat outer integument

To date, the expression patterns of *FUS3*, *NAC38,* and *NAC58* in the seed coat are unknown. Analysis of transcriptomic data shows that both *NAC* genes were preferentially expressed in the SC with stronger expression at the mature green embryo stage, where *FUS3* was also highly expressed (**Supplementary Figure 2**). To confirm these expression patterns, we generated *pNAC38:GFP* and *pNAC58:GFP* reporter lines. For both reporters, the GFP signal was present in the outer integument of seeds at the mature green stage (**Figure 2A; Supplementary Figure 3**). Transcriptomic studies also showed *FUS3* expression in the SC of torpedo/linear cotyledon and mature embryos stages, as well as embryos (**Supplementary Figure 2**). Using the *pFUS3:FUS3-GFP* and *pFUS3:FUS3ΔC-GFP* sensitive reporter, the latter lacks the PEST degradation domain, FUS3 was previously shown to be expressed in developing embryos, and the outer integument of the SC six to 48 hours after pollination (Gazzarrini et al., 2004; Lu et al., 2010; Wu et al., 2020). The *pFUS3:FUS3ΔC-GFP* sensitive reporter confirmed FUS3 localization to the outer layers of the SC at torpedo to mature green stages, in agreement with laser-microdissection studies (Le et al., 2010; Belmonte et al., 2013) (**Figure 2A, B; Supplementary Figures 2 and 3**). RT-qPCR showed expression of all three genes at 12 and 14 DAF, with *FUS3* showing the earliest and highest expression level among the three TFs (**Figure 2C)**.

**Figure 2.**
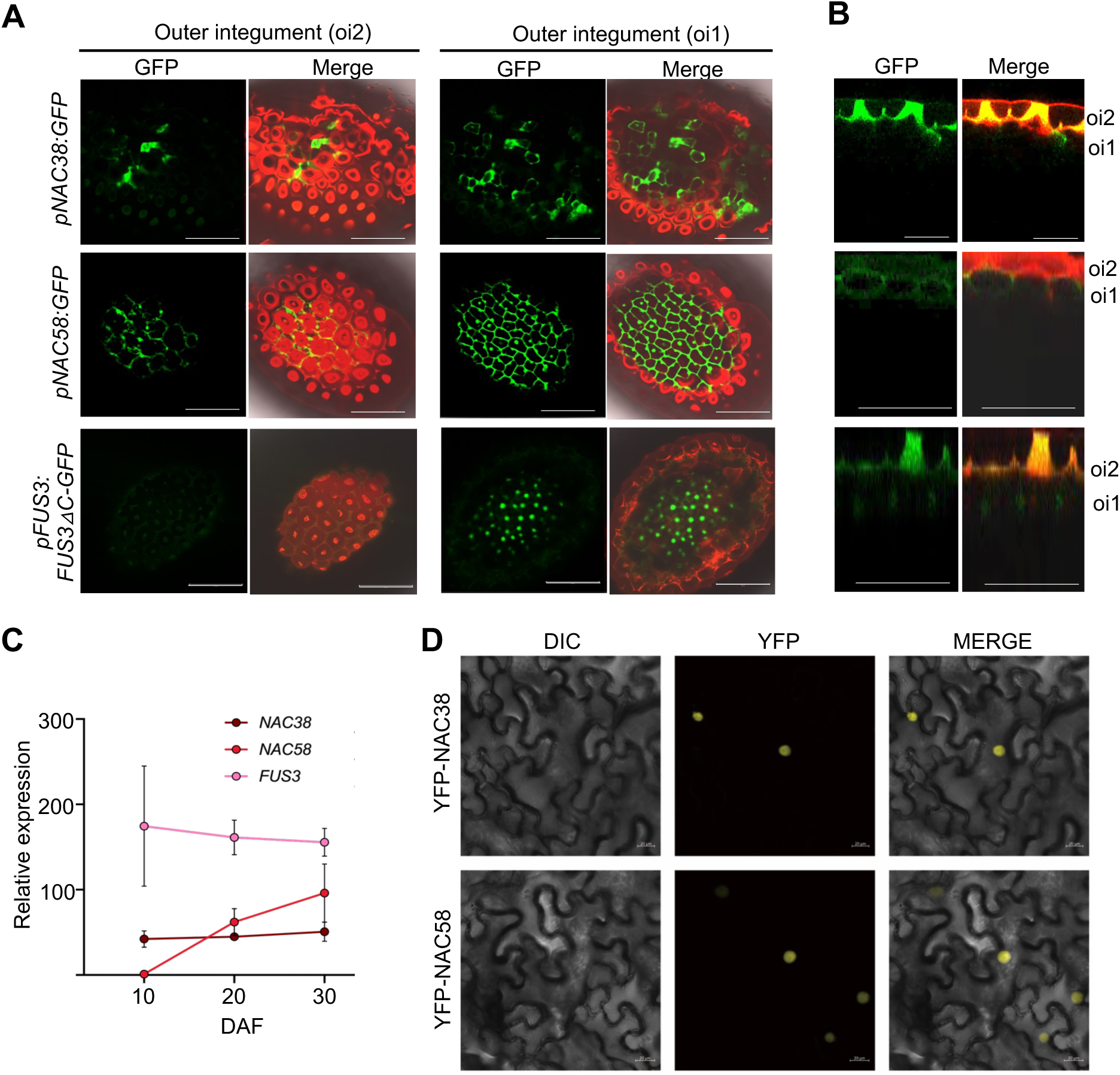
*NAC38*, *NAC58*, and *FUS3* are expressed in the seed coat outer integument. **A**, Confocal microscopy images showing GFP fluorescence of the *pNAC38:GFP*, *pNAC38:GFP*, and *pFUS3:FUS3ΔC-GFP* reporter lines in the outer integument at mature green stage. GFP and propidium iodide (PI) were excited at 488 nm. GFP emission was detected at 515-535 nm and PI at 585-735 nm. **B**, Orthogonal sections showing GFP fluorescence in the outer integument (oi) layers. Scale bars = 50 μm (A), 25 μm (B). Seeds were stained with 100 μg/mL PI. Green: GFP. Red: PI. **C**, RT-qPCR of developing seeds at 10, 12, and 14 days after flowering (DAF) showing relative *NAC38*, *NAC58*, and *FUS3* transcript levels. Averages of three biological replicates ± SEM were normalized using *ACTIN7* as the internal control. Results are plotted as the ratio to the lowest detected level set at 1.**D**, Subcellular localization of YFP-NAC38 and YFP-NAC58 in *Nicotiana benthamiana* leaves by confocal microscopy. YFP fluorescence was detected exclusively in the nuclei. YFP was excited at 488 nm and detected using a 505–530 nm band-pass filter. Scale bars = 20 µm.

We also generated NAC38-YFP and NAC58-YFP fusion proteins, which were localized to the nuclei when transiently expressed in *N. benthamiana* (**Figure 2D**). FUS3 was previously shown to be localized to the nuclei when transiently expressed in *N. benthamiana* and in stable Arabidopsis lines (Gazzarrini et al., 2004; Tsai and Gazzarrini, 2012a).

Together, these results indicate that *FUS3*, *NAC38*, and *NAC58* are co-expressed in the outer integument of the SC at the mature green stage and that the NAC transcription factors are nuclear localized.

### FUS3, NAC38, and NAC58 contribute to the differentiation of the seed coat mucilage secretory cells

To investigate the role of FUS3, NAC38, and NAC58 in SC development, we characterized the loss-of-function *fus3-3* mutant (Keith et al., 1994) and T-DNA insertion mutants, *nac38-3* and *nac58-2*, which showed strong downregulation of the respective transcripts (**Supplementary Figure 4A to C**). We also generated a *nac38-3 nac58-2* double mutant.

Given the *NACs* and *FUS3* expression patterns, we investigated their roles in the development of mucilage secretory cells in the SC epidermis. We first examined the surface of the SC through scanning electron microscopy (SEM). All mutants showed decreased radial cell wall width and *fus3-3* also showed disruption of the epidermal cell morphology indicating that FUS3, NAC38 and NAC58 promote radial cell wall thickening (**Figure 3**).

**Figure 3.**
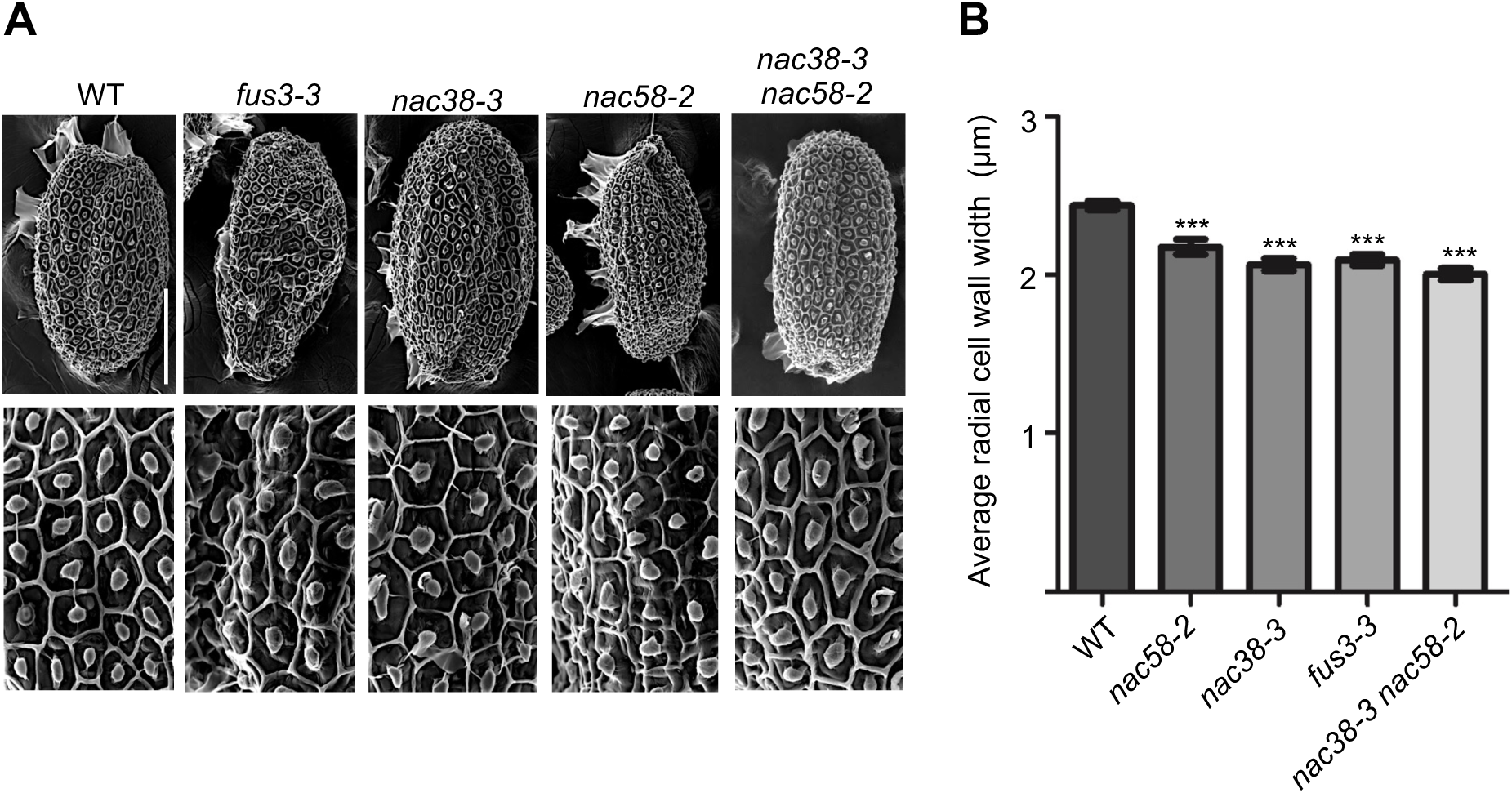
Loss of *FUS3*, *NAC38*, and *NAC58* decreases the thickness of the radial walls in the seed coat epidermal cells. **A,** Scanning electron microscopy (SEM) images of *A. thaliana* imbibed seeds of WT, *fus3-3*, *nac38-3, nac58-2,* and *nac38-3 nac58-2*. Scale bar = 200 µm in whole seed images (top) and 50 µm in surface images (bottom). **B,** Graph showing the averages of the radial cell wall width (µm) of WT and mutants. Images of 10 seeds were taken and one representative is shown. One-way ANOVA, Tukey’s post hoc test; *** p < 0.01 (N= 100 cells; 10 cells per seed, 10 seeds per genotype).

Upon imbibition, the mucilage secretory cells extrude mucilage, which produces a halo surrounding the seed (Voiniciuc et al., 2015). To test the roles of NACs and FUS3 in mucilage extrusion, we first stained the seeds with Ruthenium red, which adheres to acidic polysaccharides such as pectin in the mucilage (McFarlane et al., 2014). All genotypes were able to release mucilage, however *fus3-3* and the *nac* mutants showed decreased adherent mucilage compared to WT (**Figure 4A, top; Supplementary Figure 5A)**. EDTA chelates Ca^2+^ and reduces pectin crosslinking, thereby promoting the release of mucilage. In the presence of EDTA, all *nac* mutants released a similar amount of mucilage as WT, while *fus3-3* released a slightly larger amount than WT. Furthermore, all mutants displayed aberrant mucilage halo borders compared to WT, with *nac38*, *nac58*, and *nac38 nac58* showing spiky edges (**Figure 4A, bottom**). Last, non-adherent mucilage was released by all seed genotypes, although approximately 30% of *fus3-3* seeds showed very little non-adherent mucilage (**Figure 4B** and **Supplementary Figure 5B**). These results indicate that FUS3, NAC38, and NAC58 are required to promote mucilage extrusion and regulate mucilage capsule structure.

**Figure 4.**
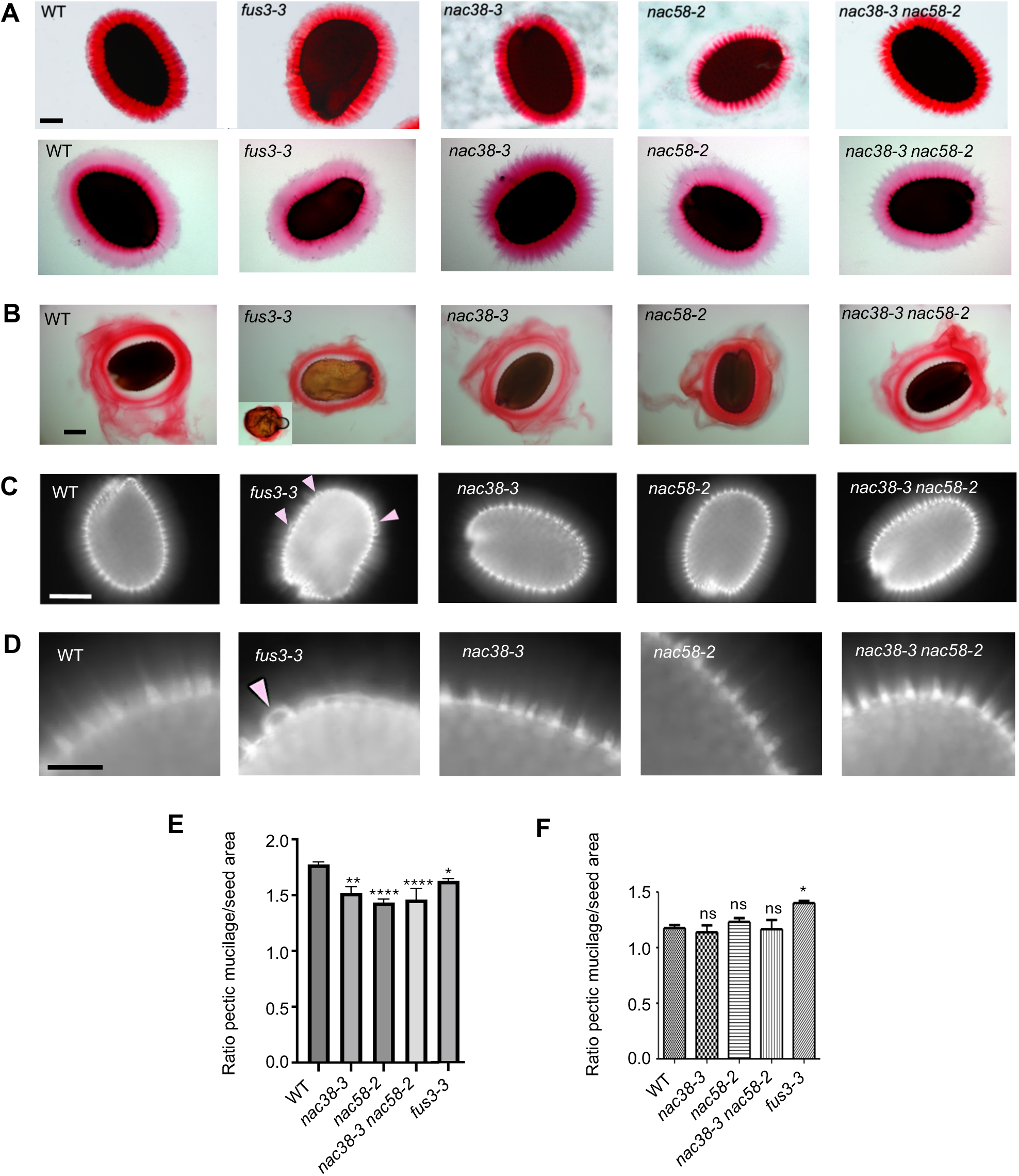
Loss of *FUS3*, *NAC38*, and *NAC58* affect seed coat mucilage extrusion and structure of the mucilage capsule. **A,** Visualization of adherent mucilage released from seeds after shacking in water (top) or water with EDTA (bottom) was stained with Ruthenium red. **B**, Non-adherent mucilage released from seeds directly stained with Ruthenium red for 30 min. **C, D,** Visualization of cellulosic rays in seeds stained with Calcofluor white. **D** Higher magnifications of images shown in **C**. Scale bars = 100 μm (A), 500 μm (B), 200 μm (C), 50 μm (D). Arrowheads indicate floppy cellulosic rays in *fus3*. **E**, Ratio of adherent mucilage area to seed area of seeds incubated with RR in water. Values represent means of three biological replicates ± SEM (n=20-30 seeds per replicate). One-way ANOVA followed by Dunnett post-hoc test (* p < 0.05, ** p < 0.01; ** p < 0.001). **F**, Ratio of adherent mucilage area to seed area of seeds incubated with RR in water + EDTA. Values represent means of three biological replicates ± SEM (n=30-40 seeds per replicate). One-way ANOVA followed by Dunnett post-hoc test (* =p < 0.05).

Next, we stained cellulose that is present in the adherent mucilage with Calcofluor. WT showed intense fluorescence in the columella and the rays stemming from it, and a diffuse fluorescence between the rays, as previously shown (Voiniciuc et al., 2015) (**Figure 4C**). In contrast, the *nac* mutants displayed a reduction of the diffused stain and/or shorter rays (**Figure 4C,D**). This pattern resembles Calcofluor staining of the *cesa5/mum3* mutant, which lacks a cellulose synthase required for mucilage cellulose production and cellulosic rays, mucilage adhesion, and radial wall thickening (**Supplementary Figure 5B)** (Sullivan et al., 2011; Mendu et al., 2011; Griffith et al., 2015). In *fus3-3*, the cellulosic rays extending from the columellae were floppy and not perpendicular to the SC surface as in WT (**Figure 4C,D**).

Collectively, these data indicate that NAC38, NAC58 and FUS3 are required for proper development of the epidermal secretory cells, including secondary radial wall thickening, mucilage capsule structure, and adherent mucilage extrusion.

### Loss of *NAC38* and *NAC58* affect seed germination and sensitivity to salinity stress after prolonged dry storage

Defects in SC epidermal cell development can compromise seed permeability, germination, and sensitivity to environmental stress. Therefore, we first tested the permeability and viability of mutant seeds with tetrazolium. Dry seeds are impermeable at maturity, however, permeable seeds can absorb the tetrazolium salt and form an insoluble, red pigment (formazan) when in contact with a viable embryo, as SC cells are dead at maturity (Chandler et al., 1997). The permeability of *fus3-3*, *nac38, nac58,* and *nac38 nac58* was similar to that of WT (**Figure 5A, B**). This was not surprising, as mucilage cellulose mutants (*cesa2 cesa5 cesa9*, *fei2, sos5, cobl2*) and mucilage pectin mutants (*mum2, mum4*) did not show increased tetrazolium stain (de Silva et al., 2021).

**Figure 5.**
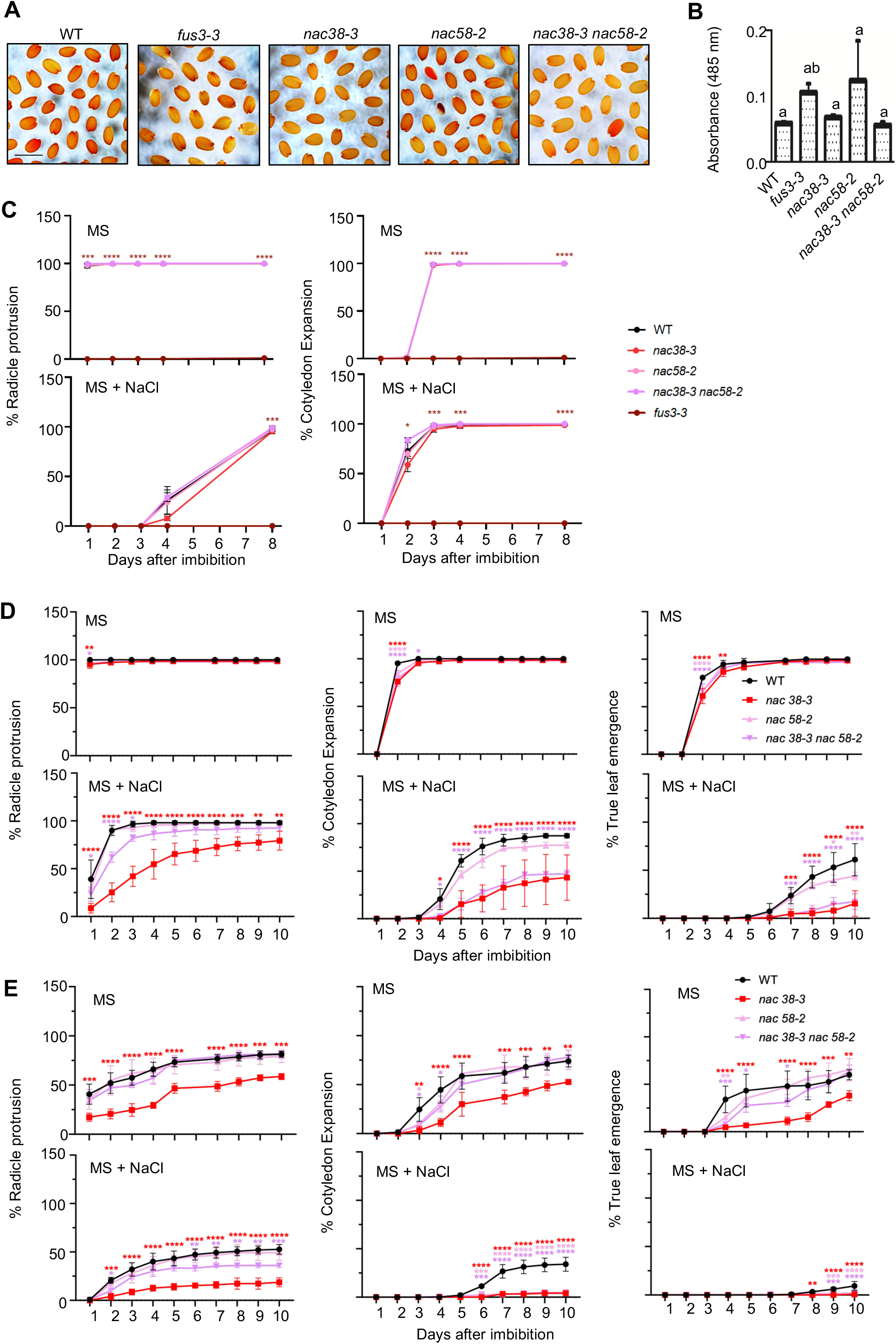
Loss of *NAC38* and *NAC58* affect seed germination and sensitivity to salinity stress. **A**, Images of WT, *fus3-3*, *nac38-3*, *nac58-2*, and *nac38-3 nac58-2* seeds incubated in 1% (w/v) Tetrazolium hydrochloride (TH). **B**, Quantification of formazans in seeds incubated with TH by absorbance at 485 nm. Average of three biological replicates ± SEM is shown (20 mg of seeds per replicate). **C-E**, Germination (radicle protrusion), seedling establishment (cotyledon expansion) and vegetative growth (true leaf emergence) rates of seeds imbibed on MS agar plates ± 150 mM NaCl. Three-months-old (**C**), 1.5-year-old (**D**), and four-years-old (**E**) seeds were surface sterilized, stratified for three days, and germinated under a long day cycle at 21°C (16h light, 8h dark). Averages of three biological replicates ± SEM. One-way ANOVA followed by Dunnett post-hoc test (* p < 0.05, ** p < 0.01; ** p < 0.001).

Next, we tested whether defects in the development of the mucilage secretory cell, in particular mucilage structure and extrusion, affected germination (radicle protrusion), seedling growth (cotyledon expansion) and vegetative growth (true leaf emergence) of freshly harvested (1 month old), and aged seeds (1.5 and 4 years old). We also tested sensitivity of the mutants to elevated salinity (150 mM NaCl), a major environmental stress for germinating seeds that can cause both osmotic and ionic/oxidative stress and can be mitigated by the mucilage (Geneve et al., 2017; Yang et al., 2021; Tsai et al., 2021). Seed germination and seedling growth rates of one-month old *nac38-3, nac58-2*, and *nac38-3 nac58-2* seeds were comparable to WT on control (MS) and NaCl (**Figure 5C**). As expected, *fus3-3* is desiccation intolerant and its germination rapidly declined after a short period of dry storage (**Figure 5C**; Keith et al., 1994).

Interestingly, after 1.5 years of dry storage, seedling and vegetative growth were slightly delayed on MS in all *nac* mutants, while sensitivity to salinity stress was strongly increased in *nac38-3* and *nac38-3 nac58-2* compared to WT, suggesting both NAC38 and NAC58 positively regulate germination during ageing, but only NAC38 appeared to regulate sensitivity to salinity stress in aged seeds (**Figure 5D)**.

Prolonged dry storage (4 years) delayed vegetative growth on MS of all genotypes, and *nac38-3* showed the stronger delay (**Figure 5E**). Furthermore, *nac38-3* showed a strong decrease of germination and vegetative growth after 10 days compared to WT, suggesting loss of *NAC38* affects seed viability (**Figure 5E**). In contrast, *nac58-2* displayed the weakest delay in vegetative growth on MS, and *nac38-3 nac58-2* partially rescued *nac38-3* decreased viability on MS, suggesting NAC38 may antagonize NAC58 in seed longevity (**Figure 5E**).

Interestingly, *nac58-2* seedlings showed a milder delay of vegetative growth on NaCl in 1.5-years-old seeds (**Figure 5D**), but an almost complete inhibition of seedling growth in 4-years-old seeds, similar to *nac38-3* and *nac38-3 nac58-2* (**Figure 5E**), suggesting NAC38 and NAC58 may work in a similar pathway to regulate seed sensitivity to salinity stress after prolonged dry storage.

These results show that NAC38 is required to promote seed longevity and antagonizes NAC58 in this process. However, both NAC38 and NAC58 regulate sensitivity of seeds to salinity stress during prolonged dry storage.

### Loss of *FUS3*, *NAC38* and *NAC58* affects the expression of genes involved in cell wall, lipid, and secondary metabolism during seed maturation

To better understand how FUS3 and the NACs regulate SC development, we conducted transcriptomic studies of *fus3-3* and *nac38 nac58* developing seeds at 12 and 14 days after flowering (DAF), when these TFs are co-expressed (**Supplementary Figure 6; Supplementary Data Set 1)**. Gene Ontology (GO) enrichment analysis of *fus3-3* showed downregulation of pathways that are known to be controlled by FUS3, such as lipid metabolism and storage, including fatty acid biosynthesis (**Supplementary Figure 7**). This agrees with the strong decrease in fatty acids shown by the *fus3-3* mutant and increase in triacylglycerol shown by ectopic *FUS3* induction during vegetative development (Roscoe et al., 2015; Zhang et al., 2016). In contrast, DNA replication, cell cycle, RNA metabolism, and ribosome biogenesis related processes were enriched among the upregulated pathways in *fus3-3* (**Supplementary Figure 7**). This is in line with ectopic activation of cell cycle genes and precocious germination of *fus3-3* immature seeds, as well as inhibition of cell cycle genes and delayed germination displayed by ectopic *FUS3* expression (Keith et al., 1994; Raz et al., 2001; Gazzarrini et al., 2004; Tsai and Gazzarrini, 2012b). Furthermore, cell wall-related processes, phenylpropanoid and secondary metabolism were also enriched among the differentially expressed genes (DEGs), including genes known to regulate mucilage biosynthesis and suberin (**Figure 6**; **Supplementary Figure 7**).

**Figure 6.**
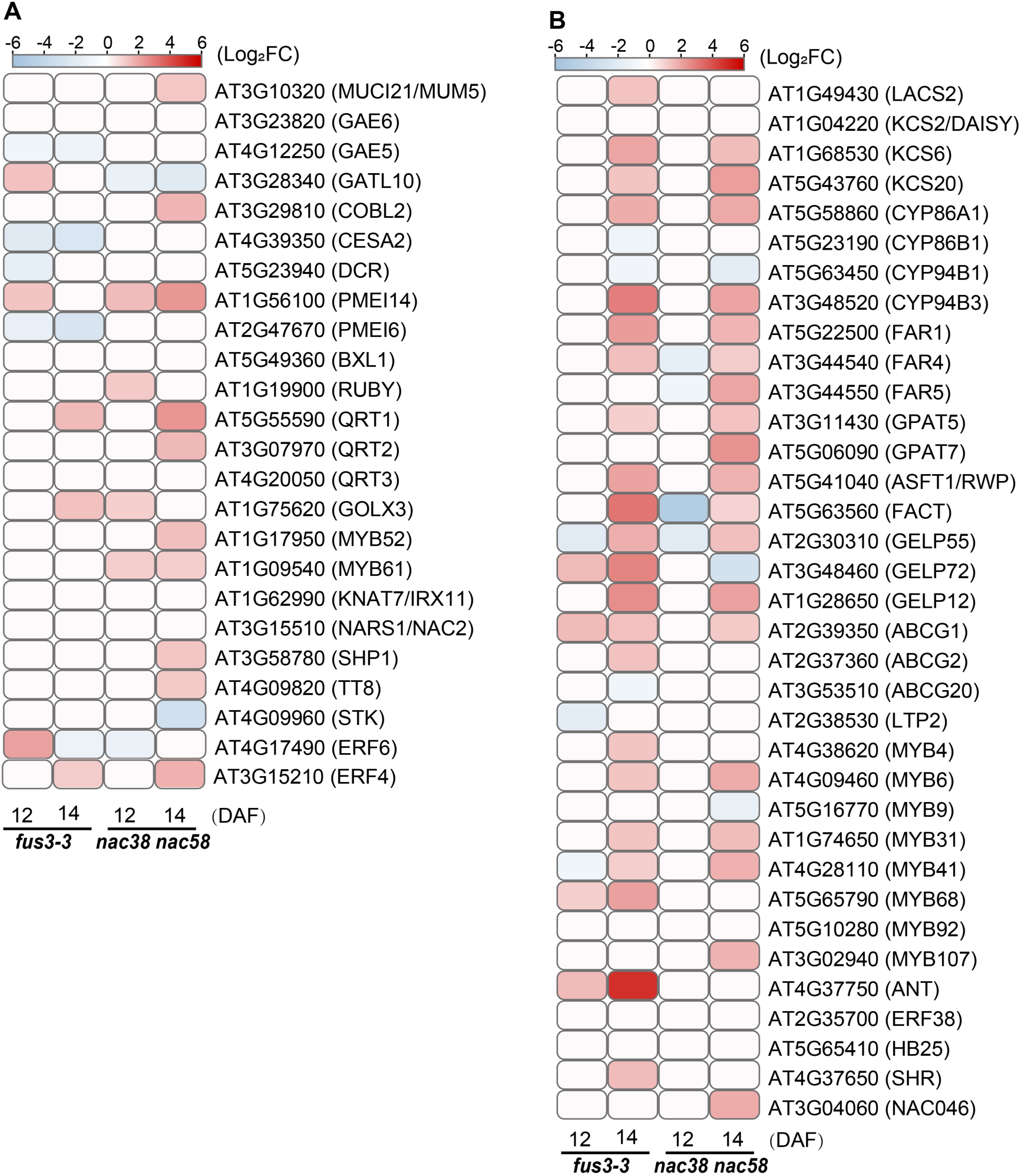
Differentially expressed genes in mucilage and suberin pathways of *fus3-3* and *nac38-3 nac58-2* seeds at 12 and 14 days after flowering. **A**, **B**, Heatmap displaying normalized expression levels (log2 fold-change relative to wild-type) of differentially expressed genes (DEGs) in mucilage (**A**) and suberin (**B**) pathways in *fus3-3* and *nac38-3 nac58-2* seeds at 12 and 14 days after flowering (DAF).

The *nac38 nac58* mutant showed very little changes in gene expression at 12 DAF compared to 14 DAF (**Supplementary Figure 7**), suggesting that *NAC38* and *NAC58* have a more prominent function during late seed coat development, in agreement with their expression patterns (**Supplementary Figure 7; Figure 2**). GO enrichment analysis showed downregulation of defense genes and differential expression of genes involved in fatty acid metabolism, phenylpropanoid biosynthesis, secondary metabolism, and nucleotide sugar metabolism (required for mucilage biosynthesis) (**Supplementary Figure 7**). The strongest enrichment was in the biosynthesis of the lipid polyesters suberin and cutin, as well as waxes, which are synthesized from fatty acid precursors and the phenylpropanoid pathway (**Figure 6**). Interestingly, suberin accumulates in the subepidermal layer (oi1), where all three TFs are co-expressed. At 14 DAF, almost half of the *nac38 nac58* DEGs were also differentially expressed in *fus3-3* (483 genes, 301 upregulated and 182 downregulated) (**Supplementary Figure 8**).

These results show that loss of *FUS3* as well as *NAC38* and *NAC58* during seed maturation alters the expression of cell wall, lipid, and secondary metabolism related genes, including mucilage and suberin, and that a subset of these genes are co-regulated by the NACs and FUS3.

### NAC58 and FUS3 regulate suberin monomer composition and FUS3 controls suberin levels in the seed

The differential expression of mucilage-related genes agrees with the observed phenotypes of *fus3-3* and *nac38 nac58* mutants (**Figures 3**, **4**). However, altered expression of genes in the suberin pathway was an interesting finding that warranted further investigation given that *NAC38*, *NAC58*, and *FUS3* are expressed in the subepidermal layer (oi1) of the SC, where suberin is deposited (**Figure 2**). The central suberin biosynthesis genes, *GPAT5*, *ASFT*, and *FAR1,4,5* are also expressed in this layer (Beisson et al., 2007; Molina et al., 2009; Domergue et al., 2010). GPAT5 is required for the accumulation of aliphatic suberin monomers, including very-long-chain fatty acids (VLCFA), ω-hydroxy fatty acids (OH-FA), and ω-dicarboxylic acids (DCA), while FAR generates the fatty alcohols through the fatty acid pathway, and ASFT incorporates ferulate, a phenylpropanoid derivative, into aliphatic suberin (**Supplementary Figure 9**) (Beisson et al., 2007; Molina et al., 2009; Domergue et al., 2010). Given that these and other suberin structural, transport, polymerization, and regulatory genes differentially expressed in *fus3* and *nac38 nac58* (**Figure 6B**), we analyzed suberin levels in these mutants and also included *nac38* and *nac58* single mutants.

The total seed-coat polyester monomer load and triacylglycerol oil content were broadly unchanged in the *nac* mutants (**Figure 7A and 7B**). However, when analyzed at the monomer level (**Figure 7C**) *nac58* and *nac38 nac58* displayed increases in certain monomer levels in parallel to decreases in others; therefore, changes were masked when the abundances of all monomers of a given class were added up. More specifically, there was: i) a strong decrease in 18:2 DCA, 16:0, 18:0 and 22:0 DCA in *nac58* and the double mutant, but an increase in 18:1 DCA in both lines; ii) lower content of 16-OH 16:0 - 22- OH 22:0 saturated fatty acids, but a marked increase in 24-OH 24:0; iii) increased ferulate and decreased coumarate in *nac58* and the double mutant; and iv) a decrease in primary alcohols. Given that most significant changes were observed in *nac58* and were not enhanced in *nac38 nac58*, these results suggest that *NAC58* may play a more prominent role in regulating suberin monomer levels in seeds, in agreement with its stronger expression in oi1 compared to *NAC38* (**Figure 2**). Given the lack of change in total suberin level in the *nac* mutants, NAC58 may act redundantly with other NACs in regulating his process.

**Figure 7.**
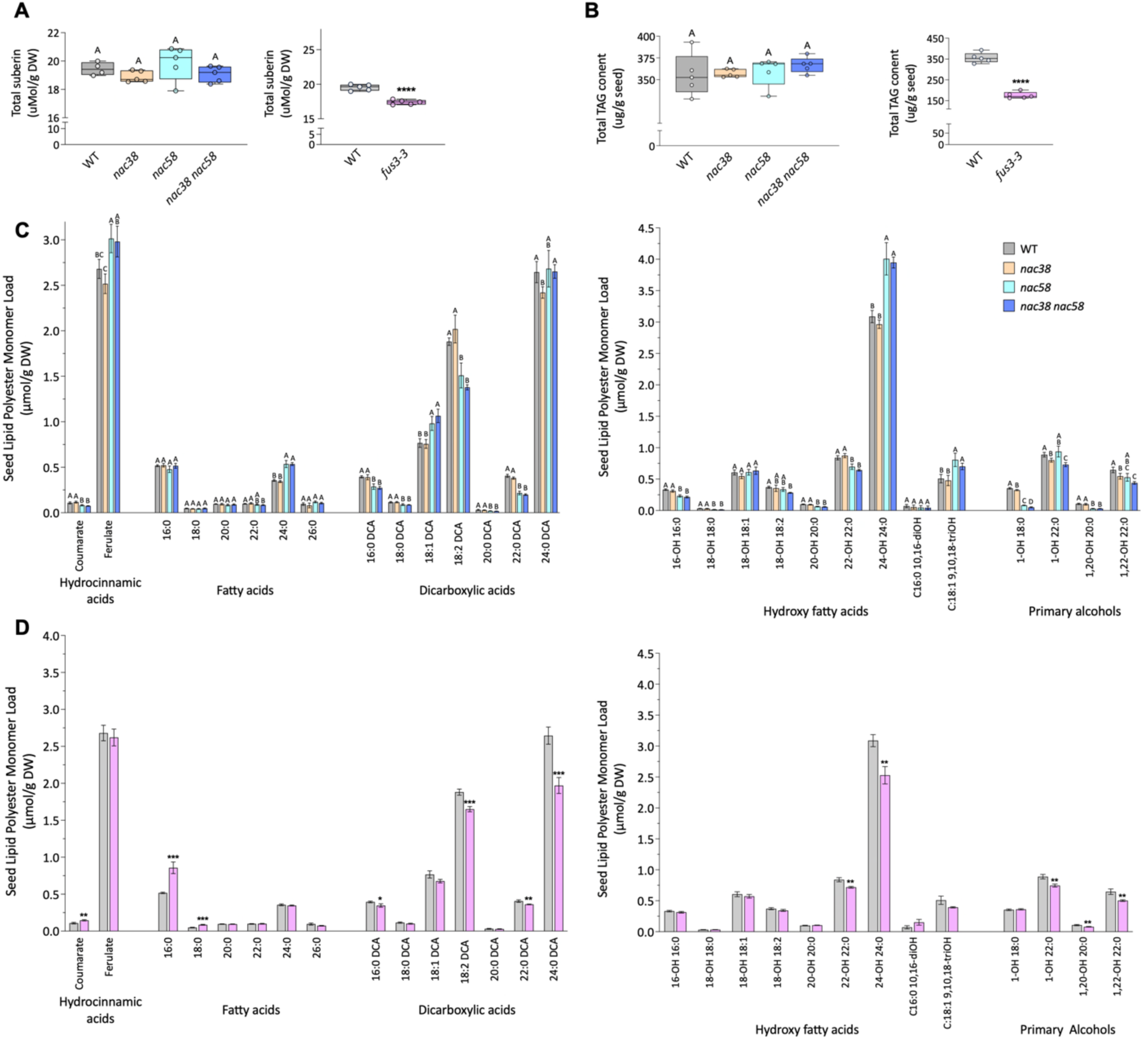
Loss of *FUS3* decreases total suberin monomers, while loss of *NAC38* and *NAC58* alters suberin monomer levels. **A**, Total seed lipid polyester monomers. **B,** Total seed triacylglycerol (TAG) content. For both: left, WT and *nac* mutants; right, WT and *fus3-3* mutant. Box-and-whisker plots show the median (center line), 25th–75th percentiles (box) and min–max (whiskers). **C, D,** Individual suberin monomer abundances in WT and *nac* mutants (**C**) and in WT and *fus3-3* mutant (**D**). Bar graphs show mean ± SD (*n* = 5) for all panels. Differences among WT and *nac* genotypes were determined by Brown–Forsythe and Welch one-way ANOVA followed by Dunnett’s T3 multiple comparisons test; different letters denote significant differences among genotypes (*P* < 0.05), and groups sharing a letter are not significantly different. Suberin monomers in *fus3-3* (pink bars) and WT (grey bars) (**D**) were compared by an unpaired two-tailed *t*-test with Welch’s correction for unequal variances. To account for multiple comparisons across the 28 monomers analyzed, *P* values were adjusted with the Holm–Šídák method (α = 0.05). Asterisks indicate significant differences from wild type: * adjusted *P* < 0.05, ** < 0.01, *** < 0.001. Total suberin and TAG load of *fus3-3* and wild type were compared separately by Welch’s *t*-test. Analyses were performed in GraphPad Prism version 11 (GraphPad Software, Boston, MA, USA).

In contrast, the loads of all main aliphatic monomers as well as total suberin were lower in *fus3-3* (**Figure 7A**, **7B**, **7C**). Oil content showed a strong decrease (∼50%) in TAGs in the *fus3-3* seeds, consistent with previous findings (∼70% to ∼80% decrease of total FA) (Tiedemann et al., 2008; Roscoe et al., 2015). These results indicate that FUS3 promotes suberin accumulation in the seed.

### FUS3 directly and indirectly regulates mucilage and suberin metabolism and regulatory genes

The transcriptomic data suggests that FUS3, NAC38 and NAC58 may regulate mucilage and suberin-related pathways important for the development of the seed coat. To find whether these TFs directly regulate these pathways, we identified their direct target genes. First we identified genes that were bound by FUS3 *in vitro* by DNA Affinity Purification (DAP; O’Malley et al., 2016) and *in vivo* by chromatin immunoprecipitation (ChIP; Wang and Perry, 2013). We then integrated the ChIP and DAP-seq, with the transcriptomic data from this study and from *FUS3* inducible lines (*iFUS3*; Yamamoto et al., 2010; **Supplementary Table 1A**). We found that many mucilage and suberin-related genes were differentially expressed in *fus3-3* and *iFUS3* lines, however, only a small subset was directly bound by FUS3. Direct targets included regulatory genes involved in mucilage (*MYB52,* the *bHLH TT8)* and suberin (*MYB41, MYB68, SHR*) regulation (**Supplementary Table 1A, C**).

These results show that FUS3 directly and indirectly regulate the transcription of genes involved in mucilage and suberin metabolism and regulation.

### Genome-wide identification of NAC38 and NAC58 binding sites *in vivo*

Next, we identified NAC38 and NAC58 targets *in vivo*. To identify genome-wide binding sites of NAC38 or NAC58, we performed chromatin immunoprecipitation sequencing (ChIP-seq) using *iNAC38* or *iNAC58* β-estradiol inducible lines (**Supplementary Figure 10**). Following β-estradiol induction, a total of ∼60.2 and ∼59.7 million clean reads were produced from *iNAC38* or *iNAC58* immunoprecipitation (IP) libraries, respectively. Approximately 39.5 million (65.54%) and 40.2 million (67.39%) reads were mapped to unique positions in the Arabidopsis (Col) genome (Genome assembly Arabidopsis version TAIR10), respectively. Additionally, 37,206 and 36,303 peaks were identified for NAC38-Input and NAC58-Input, respectively, with 11,429 and 12,576 peaks located on all chromosomes for NAC38-IP and NAC58-IP, respectively (**Figure 8A, D. Supplementary Data Set 2; Supplementary Figure 11A to C**). The genomic locations of the peaks revealed that the largest proportion of peaks (80%) for NAC38 and (78%) for NAC58 were similarly close to the transcription start site (TSS) (**Figure 8B and E).**

**Figure 8.**
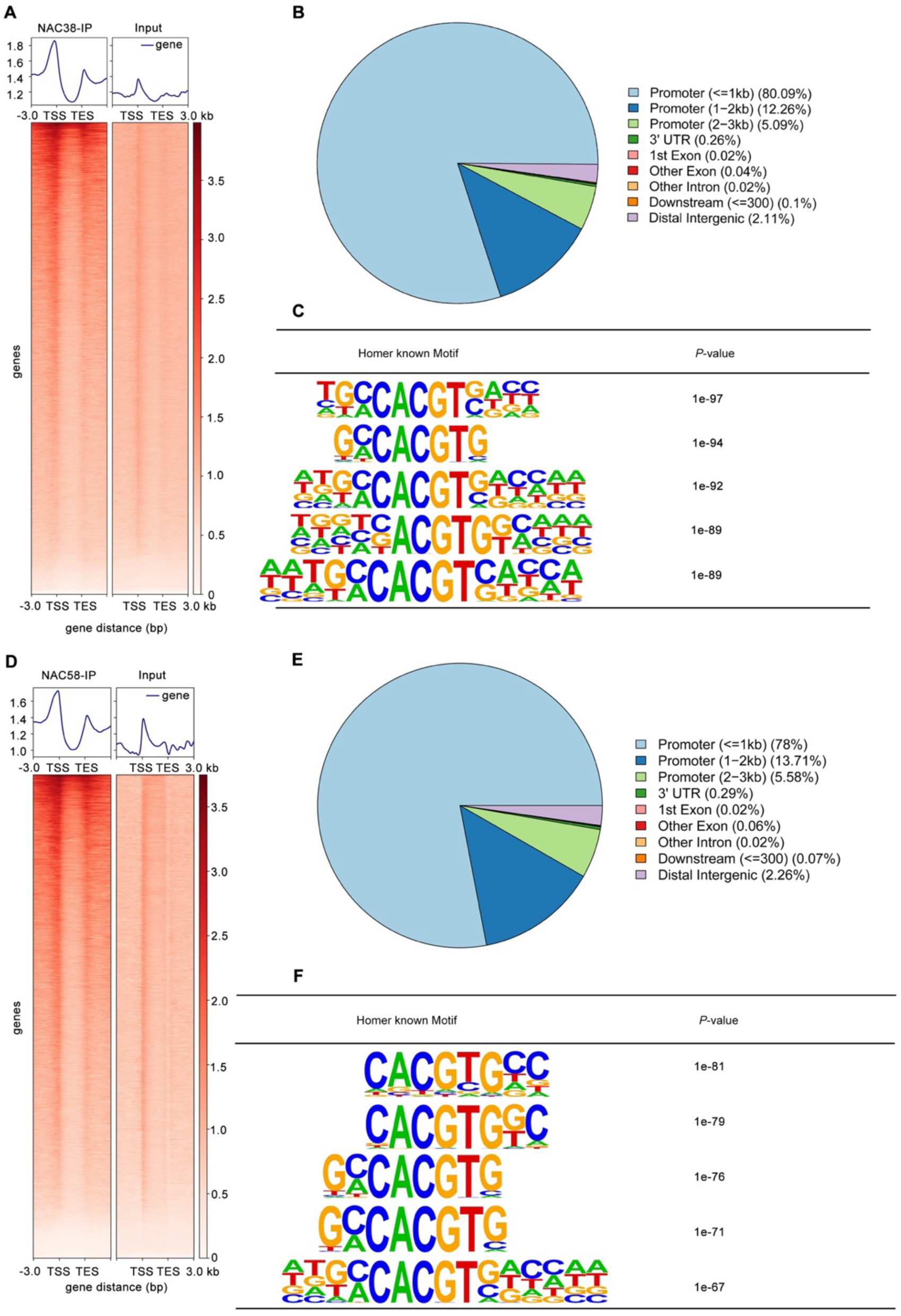
Genome-wide mapping of NAC38 and NAC58 binding sites identified by ChIP-seq. **A, D**, Heatmaps showing the occupancy profiles of NAC38 (**A**) and NAC58 (**D**) centered on transcription start sites (TSS) and transcription end sites (TES), respectively. Binding signal intensity is represented by color gradient. **B, E**, Genomic distribution of NAC38 and NAC58 ChIP-seq peaks across annotated functional regions. Functional categories are defined as follows: promoter (≤1 kb), ±1000 bp relative to the TSS; promoter (1–2 kb), 1000–2000 bp upstream or downstream of the TSS; promoter (2–3 kb), 2000–3000 bp upstream or downstream of the TSS; downstream (≤300 bp), ±300 bp relative to the TES; distal intergenic, intergenic regions located ≥1000 bp downstream of one gene’s TES and ≥3000 bp upstream of the next nearest gene’s TSS. Exons, introns, and 3′UTRs correspond to genome annotation-based gene models. **C, F**, HOMER-based enrichment analysis of known binding motifs within NAC38 (**C**) and NAC58 (**F**) peak regions. The top five most significantly enriched motifs (ranked by *p*-value) are displayed.

HOMER-based motif enrichment analysis revealed that the core NAC motif CACG[T(G/C)] was enriched for both NAC38 and NAC58 binding sites (**Figure 8C and F**). CACG was also identified as core NAC38 and NAC58 motif in DAP-seq (O’Malley et al., 2016) (**Supplementary Figure 12**). Additional binding motifs that were enriched included TTGGGCCC for NAC38 and GTGGGCCC NAC58; GGNCCC is the core consensus binding sequence of TCP TFs (**Supplementary Figure 11D and E**; Martín-Trillo and Cubas, 2010). Furthermore, the GAGA-motif was also enriched among NAC38 [(GA)_8_] NAC58 [(TC)_8_] targets and this motif is bound by BASIC PENTACISTEINE (BPC) TFs that can function as repressors and activators of gene expression (**Supplementary Figure 11D and E**) (Monfared et al., 2011; Petrella, 2020; Wu et al., 2020). Both classes of TFs are involved in reproductive development.

Collectively, these results show that NAC38 and NAC58 bind relatively similar motifs mostly in the promoters of target genes, suggesting they may regulate similar processes.

### NAC38 and NAC58 direct targets include mucilage and suberin metabolism and regulatory genes

To identify NAC38 or NAC58 direct targets (bound and differentially expressed genes), RNA-seq was performed using the *iNAC38* or *iNAC58* inducible lines. A total of 2,249 or 3,187 DEGs were found in *iNAC38* or *iNAC58* following β-estradiol treatment during germination compared with DMSO control, respectively (**Supplementary Figure 13A and B; Supplementary Data Set 3**). A comparison of the ChIP-seq and RNA-seq data revealed 962 and 1,629 direct target genes for NAC38 and NAC58, respectively (**Supplementary Figure 14A and B**). Consistent with the DEGs in seeds (**Supplementary Figure 7**), DEGs and direct targets of *iNAC38* or *iNAC58* were also enriched in the biosynthesis of secondary metabolites, phenylpropanoid and suberin biosynthesis, and nucleotide sugar metabolism (required for mucilage biosynthesis), indicating a conserved role for NAC38 and NAC58 during seed development and germination (**Supplementary Figure 14C to F**). Next, we integrated NAC38 and NAC58 direct targets identified by ChIPseq and DAP-seq datasets (O’Malley et al., 2016) and differentially expressed in *iNAC38*, *iNAC58*, and *nac38 nac58*. We found that while NAC38 and NAC58 directly regulated unique sets of genes, they also shared many common targets involved in mucilage and suberin pathways (**Supplementary Table 1B, 2D**).

Among the genes directly controlled by NAC38 or NAC58, RT-qPCR and heatmap analysis showed mucilage regulatory genes *TT8*, *MYB52, STK*, *MYB61*, *NARS1*, and *ERF4,* as well as structural genes *RUBY*, *GOXL3* and *PME16* significantly changed in expression upon *NAC38* or *NAC58* induction (**Figure 9A-C**) (Golz et al., 2018; Ding et al., 2021). Moreover, the expression levels of suberin biosynthesis and transport genes *ABCG2*, *FACT, FAR4*, *KCS2*, *ABCG2*, and *GPAT2* were up-regulated in both lines (**Figure 9A, D-E**).

**Figure 9.**
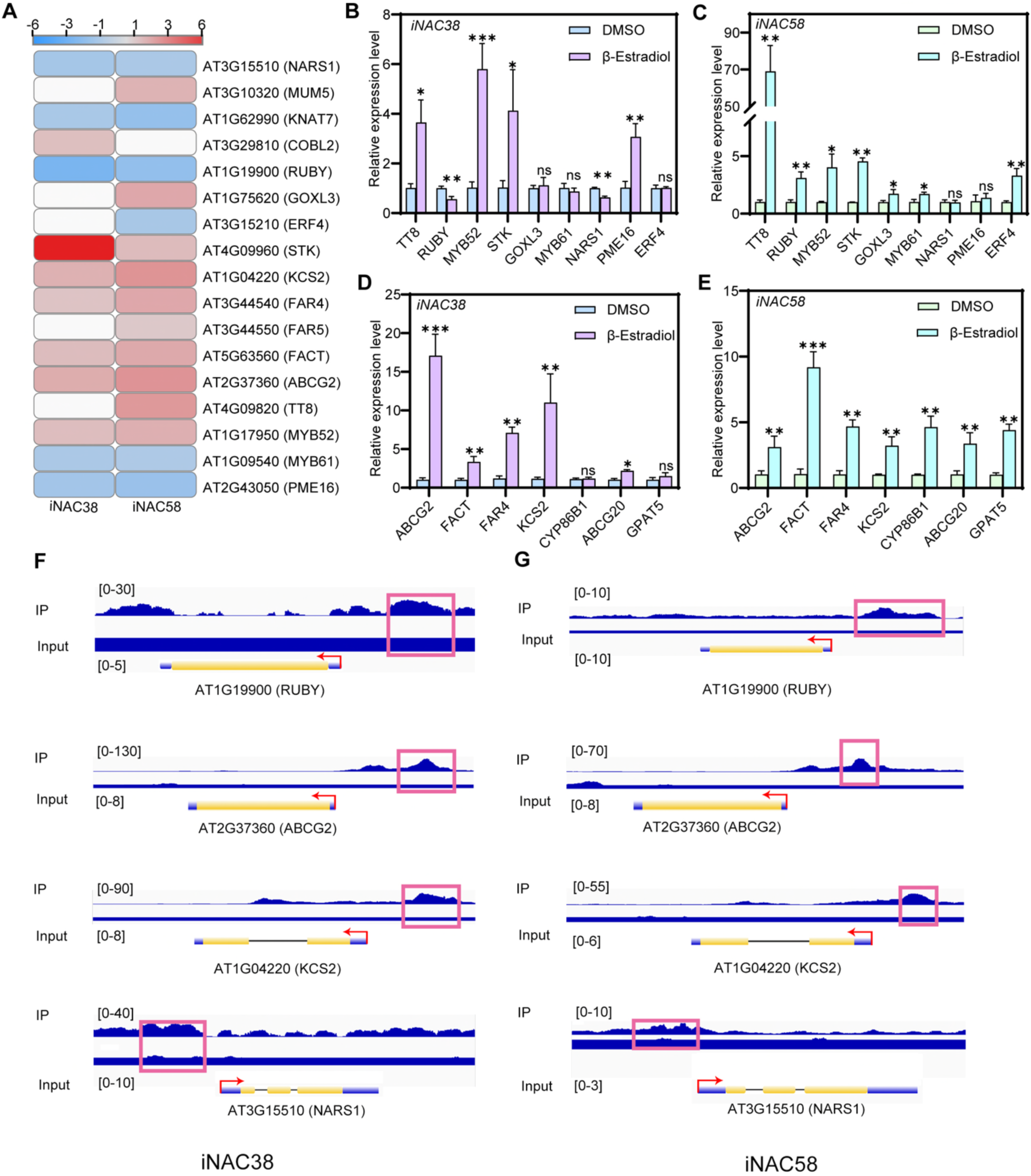
NAC38 and NAC58 associate with and directly regulate the expression mucilage and suberin pathways genes. **A**, Heatmap displaying normalized expression levels (log2 fold change relative to wild-type) of differentially expressed genes (DEGs) in mucilage or suberin pathways in *iNAC38* and *iNAC58* lines 24 h after beta-estradiol–induction. RNA-seq data are shown. **B** to **E**, RT-qPCR validation of relative transcript abundance for genes involved in mucilage (**B, C**) and suberin (**D, E**) pathways in *iNAC38* and *iNAC58* 24h after induction. Values are presented as means ± SD of three biological replicates; Student’s *t*-test (\**P* < 0.05, \*\**P* < 0.01, \*\*\**P* < 0.001). ns, no statistically significant difference. **F**, **G**, ChIP-seq binding profiles of NAC38 (**F**) and NAC58 (**G**) at select genes involved in mucilage or suberin biosynthesis, visualized using the Integrative Genomics Viewer (IGV).

Thus, our data indicate that NAC38 and NAC58 directly regulate genes involved in mucilage and suberin pathways *in vivo* and coregulate several target genes.

## DISCUSSION

During the development of the outer integument, the mucilage secretory cells undergo a dramatic differentiation process, involving radial cell wall reinforcement and the secretion of the pectin-rich mucilage, which plays a role in germination and modulate seed sensitivity to osmotic and salinity stress by unknown mechanisms (Beeckman et al., 2000; Windsor et al., 2000; Haughn and Chaudhury, 2005; Tsai et al., 2021). Meanwhile, the subepidermal layer forms an apoplastic barrier containing suberin and lignin, which confers seed impermeability and regulates dormancy and longevity (**Figure 10**) (Beisson et al., 2007; Molina et al., 2009; Yadav et al., 2014; Gou et al., 2017; Hyvärinen et al., 2025). Despite the important function of the seed coat in protecting the embryo while regulating dormancy and germination, the mechanisms and factors governing the complex development of the outer layers of the seed coat are not fully understood. Here, we show that the NAC38-NAC58-FUS3 module is required for proper development of the outer seed coat (**Figure 10**). NAC38 and NAC58 directly bound to *FUS3 in vitro* and *in vivo* and positively regulated *FUS3* expression when induced *in planta* and *in vivo*. In turn, FUS3 was shown to bind to the *NAC58* promoter *in vitro* and *in vivo* (Wang and Perry, 2013; O’Malley et al., 2016) and both *NAC38* and *NAC58* were downregulated in *fus3-3* seeds, demonstrating a feedforward positive regulatory loop between these TFs (**Figure 10**). *NAC38*, *NAC58* and *FUS3* are co-expressed in the outer integument, where they promote radial cell wall thickening and regulate mucilage capsule structure and mucilage extrusion. Furthermore, FUS3 positively regulate suberin levels, while NAC38 and NAC58 influence suberin monomers levels, although only minor changes in suberin monomer levels were found in *nac38-3*. Genome-wide binding sites and transcriptomic analysis in inducible lines showed that all three TFs directly and indirectly regulate mucilage and suberin pathway genes. Last, NAC38 plays a role in seed longevity, and both NAC38 and NAC58 modulate seed sensitivity to salt after prolonged dry storage. Collectively, our findings uncover novel regulators of mucilage and suberin pathways in the outer integument of the seed coat, which impact seed longevity and sensitivity to salinity stress.

**Figure 10.**
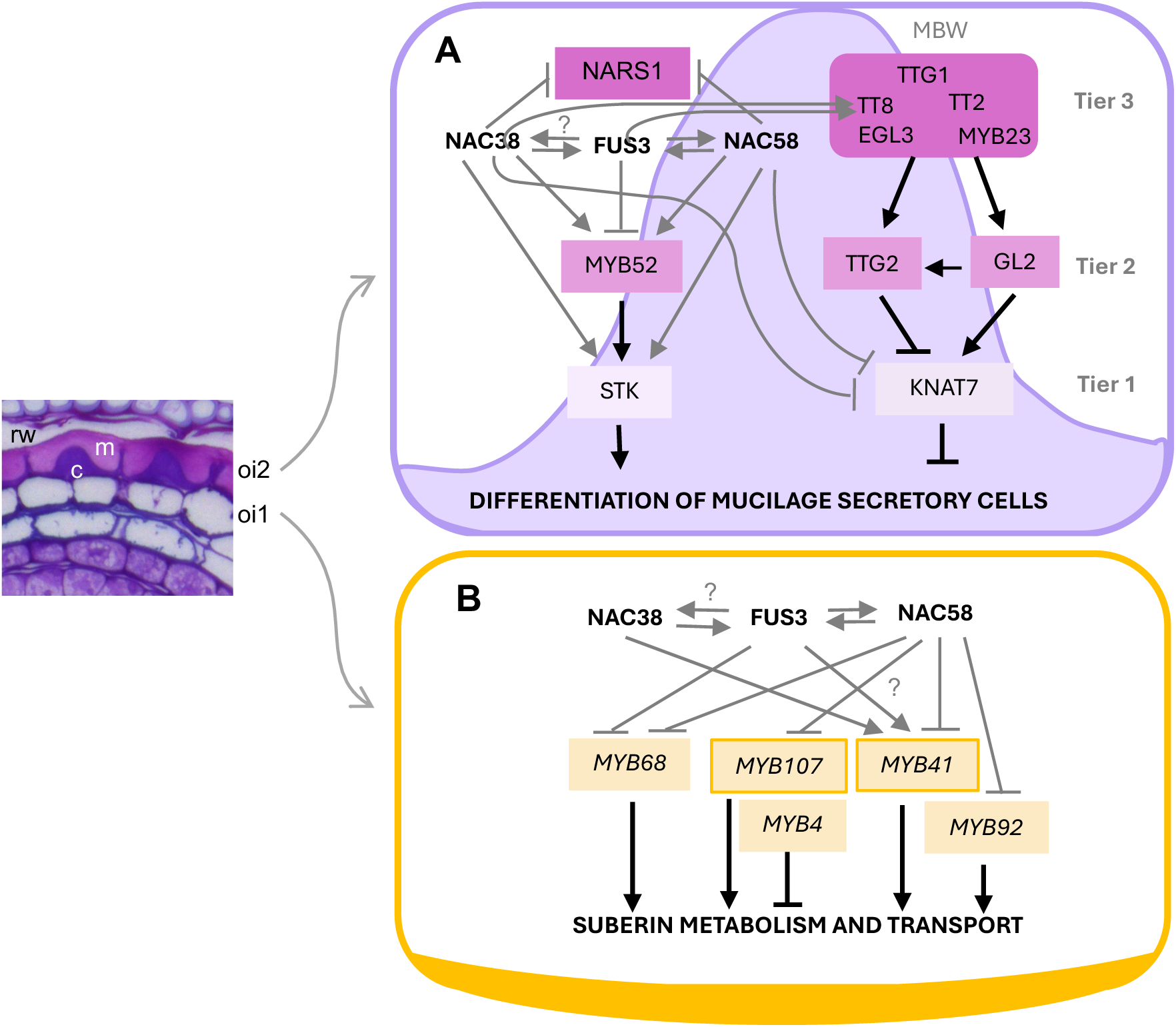
Model of FUS3-NAC38-NAC58 triad regulation of outer seed coat development. **A**, In the oi2, the MBW complex (tier 3) acts together with tier 2 (GL2 and TTG2) and tier 1 (KNAT7) transcription factors (TFs) to regulate the differentiation of mucilage secretory cells. NARS1 (tier 3), MYB52 (tier 2), and STK (tier 1) work in parallel to this pathway (Golz et al., 2018). FUS3, NAC38, and NAC58 intersects with this network of TFs at various levels, through direct binding and transcriptional regulation to regulate mucilage secretory cell development, including cell wall thickness, mucilage capsule structure, and extrusion). **B**, In the oi1, FUS3, NAC38, and NAC58 triad bind to and transcriptionally regulates MYB TFs involved in suberin metabolism and transport. FUS3 promotes suberin accumulation, while NAC38 and NAC58 regulate suberin monomer composition. MYB107 is a positive regulator of suberization in the seed, while the other MYB TFs are regulators of root suberization (Kosma et al., 2025). See text for details. Black arrows and T lines indicate previously published interactions, grey arrows and T lines are findings from this study.

### Roles of FUS3, NAC38, and NAC58 in mucilage secretory cells development

Mucilage extrusion requires the hydration and swelling of the mucilage and subsequent the rupture of the SC radial cell wall. *fus3-3* and the *nac* mutants showed thinner oi2 radial walls, reduced pectin extrusion and cellulosic rays. The mucilage extrusion defect was restored by the calcium chelator, EDTA, which facilitates mucilage extrusion by disrupting cross-links between unesterified homogalacturonan chains. This suggests *fus3-3* and the *nac* mutants have increased mucilage cohesion, possibly due to altered pectin structure or cellulose synthesis. Indeed, defects in the release of mucilage in *pmei6* mutants, which encodes a pectin methylesterase inhibitor, can be rescued by EDTA (Voiniciuc et al., 2013; Saez-Aguayo et al., 2013). Furthermore, altered cellulose synthesis in the *cesa2, cesa5* and *cesa9* mutants, which encodes cellulose synthases required for radial cell wall thickening, mucilage adhesion, and cellulosic ray formation, display similar reduction of radial wall thickening and cellulosic rays as in the *fus3* and *nac* mutants (Mendu et al., 2011; Griffith et al., 2015). Accordingly, *PMEI6*, *CESA2* and other mucilage structural and regulatory genes were differentially expressed in the *nac38-3 nac58-2* and *fus3-3* mutants, as well as in *iNAC* and *iFUS3* lines. In the SC, MYB (MYB5, TT2, MYB23), bHLH (TT8/bHLH42, EGL3/bHLH1), and WD40 (TTG1), collectively called the MBW transcriptional complex (tier 3), regulate epidermal cell differentiation together with AP2, NARS1/NAC56 and NARS2/NAC18 (**Figure 10**). They promote columella formation, thickening of the radial walls, and mucilage production and modification, in part by activating downstream targets such as *GL2* and *TTG2* (tier 2). MYB52 and MYB61 (tier 2) act in a separate pathway to activate downstream targets, *STK* and *KNAT7* (tier 1) (Francoz et al., 2015; Gou et al., 2017; Golz et al., 2018; Li et al., 2020; Allen et al., 2023; Xu et al., 2023). Our data show that the NAC38-NAC58-FUS3 module interacts with the MYB-bHLH-TTG triad at various levels (**Figure 10, Supplementary Table 1**). FUS3, NAC38, and NAC58 promote the expression of *TT8*, although direct binding to *TT8* was shown only for FUS3 and NAC38. TT2 and TT8, which are negative regulators of fatty acid accumulation in seeds, were shown to directly repress *FUS3* (Chen et al., 2014; Wang et al., 2014; Chen et al., 2015). However, it is unknown if this negative feedback regulation exists also in the seed coat. NAC38, NAC58, and FUS3 also act in parallel to the MBW complex, as they directly regulate *MYB52*, although in an opposite manner. NAC38 and NAC58 also directly bind to and repress *NARS1,* while positively regulating *STK*. Thus, NAC38, NAC58, and FUS3 coregulate shared target genes (MYB52), while also modulating the expression of different structural and regulatory genes (**Figure 10, Supplementary Table 1**). A positive feedforward regulatory loop modulates *FUS3* and *NAC58* expression, enabling a robust control of this process. *Brassica napus* ABSCISIC ACID3, BnABI3, which rescued an Arabidopsis *abi3* mutant allele (*gs1*), could directly bind to and induce the expression of *MUM4* and *GATL5* when transiently expressed in Arabidopsis (Xu and Cai, 2019). It remains to be determined whether ABI3, which belongs to the same B3 domain family as FUS3 and is also involved in seed maturation and desiccation tolerance, contributes to the regulation of mucilage levels and structure in Arabidopsis.

### Role of the FUS3-NAC module in suberin deposition

Our findings support a positive role for FUS3 in suberization of the seed, as shown by the decreased total suberin load in *fus3-3*. Furthermore, FUS3 regulates the expression of transcription factors known to promote root suberization, such as *MYB41*, *MYB52*, *MYB68*, and *SHR*, possibly implicating these TF also in seed suberization (Shukla et al., 2021b). All but *MYB41* are direct FUS3 targets (**Figure 10; Supplementary Table 1**). Interestingly, these genes were upregulated in *fus3-3* seeds, suggesting a possible feedback regulation triggered by decreased suberin levels. Of the structural genes, *FUS3* could directly regulate only a few suberin biosynthesis (*FAR1*), degradation (*GELP72*), and transport (*LTP2*) genes. Thus, FUS3 appears to directly and indirectly regulate suberin biosynthesis, possibly through MYB TFs. The decreased suberin level in *fus3-3* may also be a consequence of the decreased pool of precursors for TAGs and suberin, or decreased ABA level, since ABA is known to promote root suberization (Barberon et al., 2016; Shukla et al., 2021b) and FUS3 positively regulates ABA levels while inhibiting GA synthesis (Curaba et al., 2004; Gazzarrini et al., 2004). The reduced suberin level could contribute to the reduced dormancy and desiccation intolerance of *fus3-3* seeds (Keith et al., 1994). Indeed, previous studies have shown that suberin promotes seed dormancy and longevity. Mutants deficient in suberin monomer synthesis (*gpat5*), transport (*abcg2/6/20*) and regulation (*myb107*), have decreased seed dormancy and/or are more sensitive to seed deterioration (Beisson et al., 2007; Molina et al., 2009; Lashbrooke et al., 2016; Fedi et al., 2017; Renard et al., 2020).

The levels of suberin monomers were altered in a similar manner in *nac58* and *nac38 nac58* compared to WT, but less and different changes were observed in *nac38*, suggesting NAC58 may play a more prominent role in this process. This was surprising, given the role of NAC38 in seed longevity, and suggested that altered mucilage structure and extrusion, or other factors, played a more prominent role in *nac38* seed longevity than suberin. Nevertheless, both *iNAC38* and *iNAC58* showed upregulation of a large number of suberin biosynthesis genes, many of which were direct NAC targets (**Supplementary Table 1**). NAC58 negatively regulated TF involved in root (*MYB92, MYB68)* and seed (MYB107) suberization. Furthermore, NAC38 and NAC58 directly bound and had opposite transcriptional effects on *MYB41*, possibly explaining the lack of total suberin change in the *nac38 nac58* mutant (Gou et al., 2017; To et al., 2020; Kosma et al., 2025; Kraska et al., 2025) (**Figure 10**; **Supplementary Table 1**). Given the large family of NAC TFs (Zhu et al., 2012), NAC38 and NAC58 may act redundantly with other NACs to maintain proper suberization of the seed coat, thus explaining the unchanged total suberin level in all *nac* mutants. It is also possible that the altered structure of the mucilage secretory cells of the mutants may have triggered a compensatory response in the oi1, which resulted in the upregulation of several suberin biosynthesis and transport genes at 14 DAF. Last, a survey of the DAP-seq data indicate that many NAC TFs can associate with the *FUS3* locus. Therefore, in the absence of NAC38 and NAC58, other NACs may promote *FUS3* expression, explaining the subtle changes in suberin monomer levels and the lack of change of total suberin in the *nac* mutants.

### Role of NAC38 and NAC58 in seed longevity and salinity stress

A large body of work has elucidated key regulators of mucilage synthesis and structure; however, the physiological significance of seed mucilage remains less clear (Tsai et al., 2021). Our germination assays highlighted an important role for NAC38 and NAC58 in regulating sensitivity to salinity stress during prolonged dry storage, and a function for NAC38 in long-term seed viability in part by antagonizing NAC58. However, the mechanisms are currently unknown. Interestingly, salinity was shown to affect mucilage expansion, and mucilage has been shown to ameliorate germination under salinity stress in most but not all the species tested (Tsai et al., 2021). Furthermore, hemicelluloses have been shown to play a role in seed sensitivity to salinity stress in Arabidopsis (Yang et al., 2021). Thus, defects in mucilage structure and extrusion observed in the *nac* and *fus3-3* mutants may explain the enhanced sensitivity to salinity stress in aged seeds. Suberin has been shown to promote seed longevity (Renard et al., 2020; Renard et al., 2021). However, the lack of permeability and small changes in suberin monomers found in the *nac38-3* mutant cannot explain its decreased longevity. Although the mucilage and suberin defects combined may be responsible for the decrease in seed viability and sensitivity to salinity of the *nac* mutants during prolonged dry storage, other processes may be affected. Thus, NAC38 and NAC58 roles in sensitivity to salinity stress of aged seeds and NAC38 function in seed longevity will be the focus of future studies.

### Conclusions

We have identified two NAC domain transcription factors, NAC38 and NAC58, which bind to *FUS3,* a master regulator of seed maturation, including fatty acid biosynthesis and desiccation tolerance. NAC38, NAC58, and FUS3 are expressed in the outer integument and transcriptionally regulate genes involved in cell wall-related processes required for proper development of the outer integument, including thickening of the radial cell walls, mucilage structure and extrusion, and the biosynthesis of suberin, which are known to impact seed ageing and sensitivity to salinity stress.

## METHODS

### Plant materials and growth conditions

*Arabidopsis thaliana* Col-0 ecotype, *nac38-3* (SALK_025040.75.x) and *nac58-2* (SALK_084913) mutants were obtained from ABRC. The *fus3-3* mutant (Keith et al., 1994) and *pFUS3:FUS3△C-GFP* line (FFG△C, containing 1.5 kb of promoter upstream of transcriptional start site) (Lu et al., 2010) seeds were used for this study. Arabidopsis seeds were stored in a dry, dark place at 21°C for the indicated times. Seeds were surface sterilized in chlorine gas for 1h. Sterilized seeds were plated on ½ strength Murashige and Skoog (MS)/agar medium, stratified at 4°C for three days in the dark, and germinated in a seed incubator (Percival) under long day conditions (16h light at 22°C and 8h dark at 18°C). Radicle protrusion, cotyledon emergence, and cotyledon expansion (open at an angle of at least 180°), were scored. GraphPad Prism 10 was used for statistical analysis. Seedlings were grown on soil under long day conditions (16h light at 22°C and 8h dark at 18°C) in plant growth chambers (BioChambers).

### Generation of transgenic lines

To generate the *pNAC38:GFP* and *pNAC58:GFP* construct, a 1.5-kb upstream promoter fragments of *NAC38* or *NAC58* from the start codon site were amplified and cloned in pDONR207 entry vector by BP reaction (Invitrogen), individually. All primers are listed in **Supplementary Table 2**. The inserts were then transferred into the pGWB4 vector by LR reaction (Invitrogen). To generate the *p35S:FUS3-GFP*, *p35S:NAC38-GFP*, and *p35S:NAC58-GFP* constructs, the full-length open reading frames (ORFs) of *FUS3*, *NAC38*, and *NAC58* lacking their native stop codons were individually amplified by PCR and cloned into the pDONR207 entry vector via BP recombination (Invitrogen). Subsequently, the resulting entry clones were transferred into the destination vector pEarleyGate104 using LR recombination (Invitrogen). To generate the *NAC38* and *NAC58* inducible lines (*iNAC38* and *iNAC58*), the ORFs of *NAC38* and *NAC58*—each fused in-frame to an N-terminal HA tag—were individually amplified by PCR. The resulting amplicons were then cloned into the pER8 entry vector using *XhoI* and *SpeI* restriction digestion and ligation. Next, all constructs were transformed into *Agrobacterium tumefaciens* strain GV3101. The reporter lines were generated using the floral dip method and selected on hygromycin containing medium (Clough and Bent, 1998).

### Promoter analysis and yeast one-hybrid assay

*Cis*-regulatory element analysis of the *FUS3* promoter and genomic region was performed using PLACE (Plant *Cis*-acting Regulatory DNA Elements database; https://www.dna.affrc.go.jp/PLACE/). A 501-bp fragment spanning positions –1000 to – 501 relative to the *FUS3* transcription start site (TSS) was amplified and cloned into the pDONR-P4-P1r entry vector via BP recombination (Invitrogen). The resulting entry clone was transferred into the pDEST_His-2 bait vector using LR recombination (Invitrogen).

The bait construct was linearized with *XhoI* (New England Biolabs, NEB) and integrated into the genome of the yeast strain YM4271 (MATa) by lithium acetate (LiAc)-mediated transformation. A library of ∼1,500 Arabidopsis transcription factors (TFs) was transformed into the yeast strain EGY48 (MATα). For the initial yeast one-hybrid (Y1H) screen, diploid yeast cells co-harboring the TF library and the *FUS3* promoter bait were selected on synthetic dropout medium lacking uracil, leucine, and histidine (SD–Ura– Leu–His), supplemented with 5 mM 3-amino-1,2,4-triazole (3-AT; BioShop). Plasmids isolated from His+ colonies were Sanger-sequenced to identify interacting TFs. To validate direct binding of NAC38 or NAC58 to the *FUS3* promoter, pairwise mating was performed between EGY48 (MATα) strains expressing full-length NAC38 or NAC58 (in pGADT7) and YM4271 (MATa) strains harboring the *FUS3* promoter bait. Diploids were selected on SD–Ura–Leu–His + 5 mM 3-AT. To test binding to the *FUS3* genomic region located 1,125–1,851 bp downstream of the TSS, the Y1HGold–GAL4–AbA reporter system (Clontech) was employed. This genomic fragment was amplified and cloned into the pBait-AbAi vector using *SalI* and *HindIII* restriction sites. Because full-length NAC38 or NAC58 expressed from pGADT7 impaired growth of Y1HGold, truncated versions— NAC38 (1–171 aa) and NAC58 (1–200 aa)—were amplified and subcloned into pGADT7 using *EcoRI* and *BamHI*. The pBait-AbAi–FUS3 construct was linearized with *BstBI* (NEB) and integrated into the Y1HGold (MATa) genome via LiAc transformation. Subsequently, pGADT7–NAC38 (1–171 aa) or pGADT7–NAC58 (1–200 aa) was transformed into the Y1HGold strain stably harboring the integrated pBait-AbAi–FUS3 (1125–1851 bp) bait. Protein–DNA interactions were confirmed by growth selection on SD–Ura–Leu medium containing 300 ng/mL aureobasidin A (AbA).

### Dual-luciferase reporter assay

The *FUS3* promoter fragment (–1000 to –501 bp relative to the TSS) or the *FUS3* genomic region (1125–1851 bp downstream of the TSS) was cloned into the pGreenII 0800-LUC vector using *KpnI* and *NcoI* restriction sites (Takara Bio). The full-length open reading frames (ORFs) of *NAC38* and *NAC58* were individually cloned into the pCAMBIA1300 vector using *XbaI* and *BamHI* restriction sites (Takara Bio). Recombinant plasmids were introduced into *Agrobacterium tumefaciens* strain GV3101 by electroporation. For transient expression assays, equal volumes of Agrobacterium cultures (OD600 = 0.8) carrying the reporter (pGreenII 0800-LUC derivatives) and effector (pCAMBIA1300-NAC38 or pCAMBIA1300-NAC58) constructs were mixed and co-infiltrated into the abaxial surface of fully expanded leaves of 6-week-old *Nicotiana benthamiana* plants. Luciferase activity was measured 48 h post-infiltration using the Dual-Luciferase Reporter Assay System (Yeasen Biotechnology, Shanghai, China), following the manufacturer’s protocol. Firefly luciferase (LUC) activity was normalized to Renilla luciferase (REN) activity (internal control), and results are presented as the LUC/REN ratio.

### Scanning electron microscopy (SEM)

To view the seed surface of dry mature seeds, the seeds were imbibed in ddH_2_O for 1 hour and dried at 37°C overnight. Seeds were then coated with 40nm of osmium tetroxide (OsO_4_) and viewed with the Hitachi S530-SEM microscope. Radial cell wall width was measured using ImageJ. Four measurements per cell were collected for ten cells per image.

### RNA isolation and transcript analysis

For seeds RNA isolation and Real-Time quantitative PCR (RT-qPCR), total RNA was extracted from 10, 12 and 14 DAF seeds isolated from *A. thaliana* siliques. Ten siliques per sample were opened with a needle along the replum and seeds were gently removed and collected in a microfuge tube, flash-frozen in liquid N_2_, and stored at –80°C. Three biological replicates were prepared for each genotype at each time point. Total RNA was isolated using a RNeasy plant mini kit (Qiagen). Seeds were crushed in liquid N_2_ using micro pestles. For RNA extraction from seeds, RLC buffer was used. cDNA was synthesized using the SensiFAST cDNA synthesis kit (BioLine). RT-qPCR reactions were performed using the Quant Studio 3 Real-Time PCR system (Applied Biosystems) with SYBR green (Wisent Bioproducts). The relative expression level was calculated with the 2^-ΔΔCt^ method. *ACTIN7* and *PP2AA3* were used as the internal reference genes (Lu et al., 2010; Tsai and Gazzarrini, 2012a).

For seedling RNA isolation (*iNAC*) and RT-qPCR, total RNA was extracted from 5-day-old Arabidopsis seedlings using the FastPure® Plant Total RNA Isolation Kit (Polysaccharides & Polyphenolics-rich) (Cat. RC401-01; Vazyme, Nanjing, China), following the manufacturer’s instructions. First-strand cDNA was synthesized from 1 µg of total RNA using the HiScript® III 1st Strand cDNA Synthesis Kit (+ gDNA Wiper) (Cat. R312-01; Vazyme, Nanjing, China). Quantitative real-time PCR was performed on a Bio-Rad CFX Connect Real-Time System using iTaq Universal SYBR® Green Supermix (Cat. 1725124; Bio-Rad, Hercules, CA, USA). Thermal cycling conditions consisted of an initial denaturation at 95 °C for 3 min, followed by 40 cycles of 95 °C for 15 s, 60 °C for 30 s, and 72 °C for 45 s, with a final melting curve analysis. *ACTIN7* was used as the reference gene.

### RNA-seq and data processing

For the Identification of differentially expressed genes (DEGs) in seeds of *nac38-3 nac58-2* and *fus3-3*, RNA-sequencing (RNA-seq) was conducted at the University of Toronto Centre for the Analysis of Genome Evolution & Function (CAGEF). The quality and quantity of the total RNAs were assessed using the RNA Nano kit for the Bioanalyzer 2100 (Agilent Technologies, Palo Alto, CA). The NEBNext Poly(A) mRNA Magnetic Isolation Module (New England Biolabs, Ipswich, MA, USA) was used to select for mRNA. Sequencing libraries were prepared using the MGIEasy RNA Directional Library Prep protocol (MGI Americas, San Jose, CA, USA) following the recommended protocol. The final library was sequenced on the MGIDNBSEQ G400RS sequencer (MGI Americas, San Jose, CA), according to the manufacturer’s instructions, using the 300-cycle sequencing kit and generating 150bp X 2 paired-end reads. RNA-seq data analysis was conducted in the package systemPipeR (Backman and Girke, 2016), with limited modifications. The function trimLRPatterns from the Biostrings package was applied to trim adaptors. The short read aligner Hisat2 (Kim et al., 2015) was used to align the reads to the Arabidopsis genome. The read mapped to gene exons were counted with the function summarizeOverlaps (Lawrence et al., 2013). Over 91 % of the reads were uniquely mapped to the TAIR10 Arabidopsis genome, resulting in 82-196 million mapped reads per sample. The edgeR package (Robinson et al., 2010) was used for DEGs analysis with the glm method. The DEGs were filtered with an FDR less than 0.05 and fold change greater than 2. The DEGs overlap among the sample pairs was computed with the function overLapper and a Venn diagram was plotted with the function VennPlot. The heat map of DEGs was drawn using Pheatmap (https://cran.r-project.org/web/packages/pheatmap/).

To identify DEGs in 5-d-old seedling of *NAC38* and *NAC58* inducible lines (*iNAC38* and *iNAC58*), the seeds of *iNAC38* and *iNAC58* were surface sterilized on a shaker for 7 minutes in a 25% bleach (1.5% hypochlorite, v/v) solution, followed by five rinses with ddH_2_O. Sterilized seeds were grown on ½ strength Murashige and Skoog (MS) medium and stratified at 4°C for 3 days in the dark. 5 days after germination, seedlings were treated with either 10 μM β-estradiol or an equivalent volume of DMSO for 24 hours. Seedlings were harvested into 1.5-mL microcentrifuge tubes, immediately flash-frozen in liquid nitrogen, and stored at –80°C until RNA extraction and sequencing. Three independent biological replicates were collected per condition.

RNA sequencing was performed by Frasergen (Wuhan, China) using the DNBSEQ-T7 platform. Paired-end sequencing was conducted with a read length of 150 bp. Raw reads were subjected to quality control and preprocessing using SOAPnuke, which removed reads containing adapter sequences, reads with >10% N bases, reads consisting exclusively of poly-A tails, and reads in which >50% of bases had a quality score (Q) ≤ 20. To deplete ribosomal RNA (rRNA), a custom rRNA reference database was constructed by extracting all rRNA sequences annotated in the NCBI GenBank database using rRNA-associated keywords. Clean, non-rRNA reads were then aligned to the TAIR10 reference genome of Arabidopsis thaliana using HISAT2. Transcript assembly and abundance estimation were performed with StringTie using the HISAT2 alignment outputs; gene expression levels were quantified as fragments per kilobase of transcript per million mapped fragments (FPKM). Differential expression analysis was carried out using DESeq2 (v1.20.0) in *R* (v3.6.0) via RStudio. Read counts were normalized using DESeq2’s median-of-ratios method, and statistical testing was performed to compute Wald *p*-values. False discovery rate (FDR)-adjusted *p*-values (Benjamini–Hochberg correction) were used to control for multiple testing. Genes with an FDR-adjusted p-value < 0.05 and |log2 fold change| > 1 were designated as differentially expressed genes (DEGs).

### ChIP-seq

ChIP assays were performed on 5-day-old seedlings (harvested concurrently with seedling RNA-seq samples) expressing *iNAC38* or *iNAC58* and treated with 10 μM β-estradiol or an equivalent volume of DMSO for 24 h. The procedure was carried out by Shandong Xiuyue Biotechnology Co., Ltd. Crosslinking was performed by vacuum-infiltrating seedlings in 1% (w/v) formaldehyde for 10 min, followed by quenching with 0.125 M glycine for 5 min. Chromatin was isolated and sheared to fragments of ∼200– 500 bp using a Bioruptor® Pico sonicator (30 s on/30 s off, 27 cycles). Prior to immunoprecipitation, 20% of the sonicated chromatin was reserved as input control (designated NAC38-Input or NAC58-Input); the remaining 80% was subjected to IP using rabbit polyclonal anti-HA antibody (clone C29F4; Cell Signaling Technology, Danvers, MA, USA; Cat. #3724). Immunoprecipitated chromatin (NAC38-IP or NAC58-IP) and input samples were reverse-crosslinked, and DNA was purified. Sequencing libraries were constructed from ≥10 ng of immunopurified or input DNA using the KAPA HyperPrep Kit (Roche), and sequenced on an Illumina NovaSeq 6000 system in paired-end mode (2 × 150 bp). Raw reads were quality-trimmed and filtered using FASTX-Toolkit (v0.0.14), retaining only reads with Phred quality scores ≥Q20 across ≥90% of bases. Clean reads were aligned to the TAIR10.1 reference genome (*Arabidopsis thaliana*) using Bowtie2 (v2.3.5.1). Duplicate reads were removed using Picard MarkDuplicates (v2.1.0), and only uniquely mapped, non-duplicate reads were retained for downstream analysis. Peak calling was performed using MACS2 (v2.1.0) with default parameters against respective input controls. Peaks were annotated to genomic features using ChIPseeker (v1.20.0). For motif enrichment analysis, AME (Analysis of Motif Enrichment) from the MEME Suite was applied using the same background model (shuffled input sequences) and negative control (input DNA) as used in peak calling; enriched motifs were reported at an *E*-value threshold ≤10 (Fisher’s exact test). The JASPAR2022_CORE_plants_non-redundant database (v2) was used as the motif reference. Genome-wide signal visualization was performed using the Integrative Genomics Viewer. The sequencing data have been deposited in the National Center for Biotechnology Information (NCBI) BioProject database under accession number PRJNA1227269 and are publicly accessible as of February 24, 2025.

### Ruthenium red staining

Mature dry seeds were imbibed in 50 mM EDTA solution made with doubly-distilled water (ddH_2_O) and subjected to vigorous shaking for 1 hour. The EDTA solution was removed and stained with 0.01% (w/v) ruthenium red in ddH_2_O (Sigma-Aldrich; Cat: R275-1) by vigorous shaking for 1 hour. Images for mucilage calculations were taken using the Nikon SMZ1500 dissection microscope. Images were processed for visible mucilage using ImageJ and an adapted macro program called PIDIQ (Laflamme et al., 2016). Representative images were taken using the Zeiss Epi-fluorescence microscope.

### Tetrazolium salt assay

Dry mature seeds (15 mg) that were 6-12 weeks old were incubated in 500 ul of 1% (w/v) 2, 3, 5-Triphenyltetrazolium chloride (Sigma, Cat: T8877-10G) for 96 hours at 30°C in the dark. After incubation, samples were rinsed twice with ddH_2_O and seeds were resuspended in 1 mL of 95% ethanol. The seeds were finely crushed with mortar and pestle and the final volume was adjusted to 2 mL with 95% ethanol with the ground seeds. Tubes were centrifuged at 15 000 g for 3 minutes and 1 mL of supernatant was measured for absorbance at 485 nm. Seeds were also imaged before crushing using the Nikon SMZ1500 dissection microscope.

### Calcofluor white assay

10-20 dry mature seeds were incubated in 1 mL of ddH_2_O with mild agitation for 2 hours at room temperature. After incubation in ddH_2_O, remove seeds with 40uL of the water into depression well slides (Diversified Biotech, Cat: MIWE-2000). 5uL of 10% KOH (w/v) was added to the well, mixed by pipetting, and incubated for 2 minutes. 10uL of Calcofluor White stain (Sigma-Aldrich, Cat: 18909-100ML-F) was added to the well, mixed by pipetting, and incubated for 1 minute. Seeds were imaged using the Zeiss Axioplan 2 Epifluorescent Upright Microscope under UV light settings.

### Seed coat lipid polyesters

To determine seed coat lipid polyesters, 100 mg dry seeds were used per replicate (5 replicates per genotype), and followed published methods (Molina et al., 2006; Jenkin and Molina, 2015), with an additional solvent extraction step to ensure thorough removal of seed oils. Internal standards (50 ug each) were methyl heptadecanoate, pentadecanolactone and pentadecanol. Seed oil was analyzed according to (Li et al., 2006). Briefly, dried seeds (10 mg) were placed in a glass tube with Teflon-lined screw cap and 1 ml of 5% (v/v) conc. sulfuric acid in MeOH, 100 ug of triheptadecanoin (triacylglycerol internal standard) and 300 ul of toluene as co-solvent. The mixture was incubated at 85 C for 90 min. After cooling to room temperature, fatty acid methyl esters (FAMEs) were extracted with 1V hexane. After transferring to a new tube, the hexane phase was evaporated under nitrogen. Samples were redissolved in 50 ul hexane and run by GC-FID on a FAME column, following the chromatographic method described by (Li et al., 2006).

### Accession numbers

Sequence data from this article can be found in the Arabidopsis Genome Initiative or GenBank/EMBL databases under the following accession numbers: FUSCA3 (At3g26790), ANAC38 (AT2G24430), ANAC58 (AT3G18400).

## SUPPLEMENTARY DATA

**Supplementary Figure 1.** Sequence analysis of Arabidopsis *FUS3*, *NAC38* and *NAC58*.

**Supplementary Figure 2.** Expression profiles of *FUS3*, *NAC38*, and *NAC58* in Arabidopsis.

**Supplementary Figure 3.** Expression patterns of *FUS3*, *NAC38* and *NAC58* in developing seeds.

**Supplementary Figure 4.** Isolation and molecular characterization of *nac38, nac58*, and *nac38 nac58* mutants.

**Supplementary Figure 5.** Cellulose and pectin staining in dry seeds.

**Supplementary Figure 6.** Differentially expressed genes at 12 DAF and 14 DAF in *fus3-3* and *nac38 nac58* by RNA-seq.

**Supplementary Figure 7.** GO biological function enrichment analysis in differentially expressed genes at 12 and 14 DAF in *fus3-3* and *nac38 nac58*.

**Supplementary Figure 8.** Overlap in differentially expressed genes at 12 and 14 DAF between *fus3-3* and *nac38 nac58*

**Supplementary Figure 9.** Suberin metabolism pathways

**Supplementary Figure 10.** Generation of *iNAC38* and *iNAC58* inducible lines

**Supplementary Figure 11.** ChIP-seq characterization of NAC38 and NAC58 binding landscapes.

**Supplementary Figure 12.** Motifs bound by NAC38 and NAC58 in DAP-seq

**Supplementary Figure 13.** Identification of DEGs in *iNAC38* and *iNAC58* inducible lines.

**Supplementary Figure 14.** Identification of NAC38 and NAC58 direct target genes (bound and differentially expressed).

**Supplementary Table 1**. Suberin and mucilage related genes that are differentially expressed in *fus3-3*, *nac38 nac58,* and *iNAC* lines, and bound by FUS3, NAC38 and NAC58.

**Supplementary Table 2**. List of primers used in this study.

**Supplementary Data Set 1**. RNA-seq of WT, *fus3-3,* and *nac38 nac58*

**Supplementary Data Set 2**. ChIP-seq of *iNAC38* and *iNAC58*

**Supplementary Data Set 3**. RNA-seq of *iNAC38* and *iNAC58*.

## ACKNOWLEDGEMENTS

We would like to thank the Centre for the Analysis of Genome Evolution & Function (CAGEF) and Yunchen Gong for the bioinformatic analysis of the RNA-seq raw data. This work was supported by the Natural Science and Engineering Research Council (NSERC) of Canada (Discovery Grant 480529) to S.G. and Fundamental Research Funds for the Central Universities (SWU-KT26041 & SWU-KR22042) to L.W.

## AUTHOR CONTRIBUTIONS

Lin Wu and Jasmin Patel conducted most of the experiments; Lin Wu, Jasmin Patel, and Sonia Gazzarrini conceived the study and wrote the article with input from other authors; Myles Matundan contributed to the RNA-seq in seeds and data analysis; Maria Barsamyan performed Ruthenium red staining; Richard Yang conducted germination assays; Yang Yang performed Dual-luciferase reporter assay; Yunling Ye did ChIP-seq and Dianqiu Lv performed RNA-seq in iNAC lines; Jian Wu conducted yeast one-hybrid assays; Marc Mohammadi and Isabel Molina, extracted and quantified suberin monomers; all authors read and approved the article. We thank several undergraduate students who helped selecting and genotyping mutants and transgenic lines and conducting preliminary germination assays (Kasra Rezaei, Sebastian Emmanuel, Tiffany Tse, Nathalie Wong, Quiyun Shao).

