## Supplementary Figures for "The NAC38-NAC58-FUSCA3 module regulates outer integument differentiation in the seed coat of *Arabidopsis thaliana*"

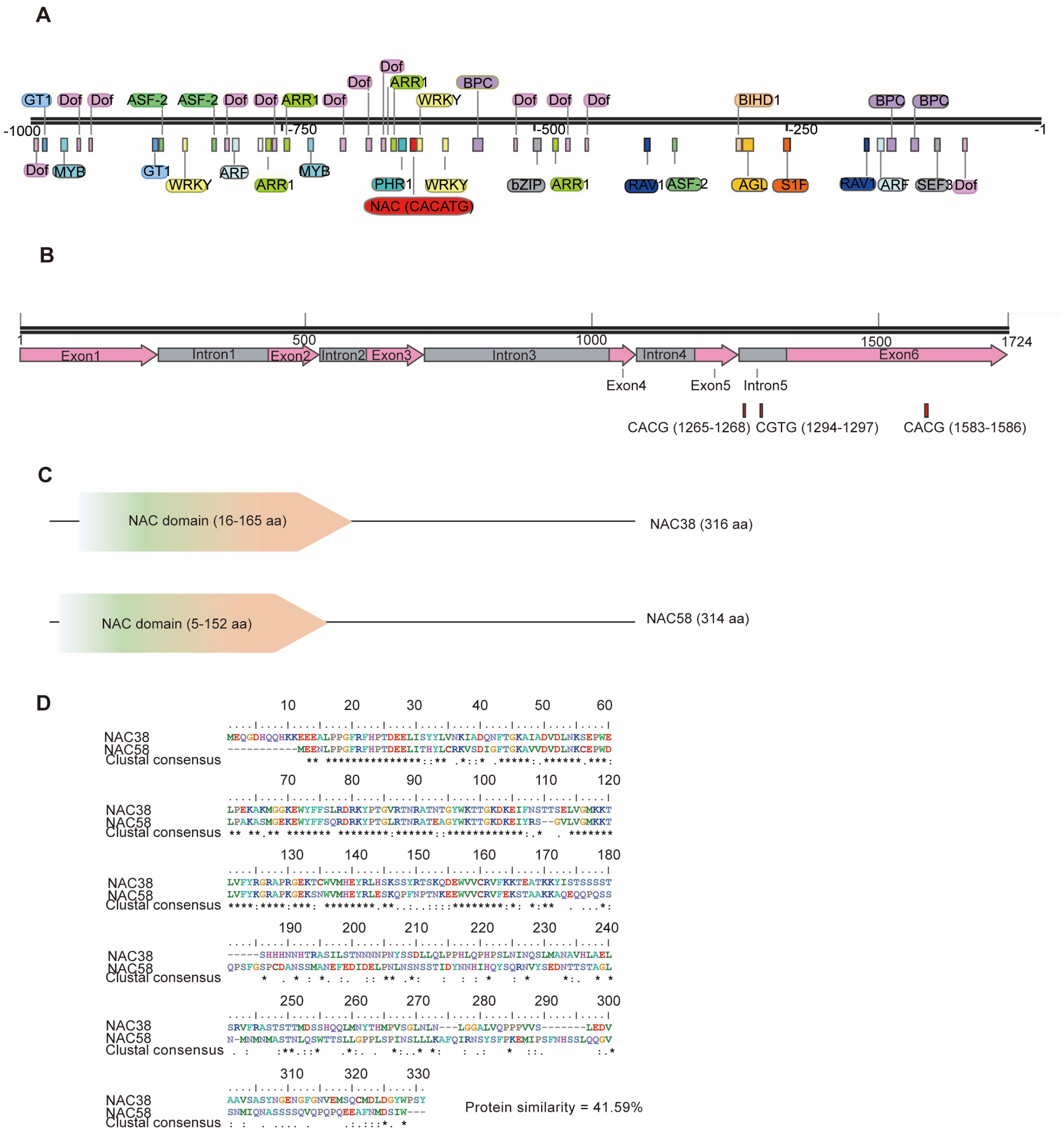


**Supplementary Figure 1. Sequence analysis of Arabidopsis *FUS3*, *NAC38* and *NAC58*.**
**A**, Schematic representation of the *FUS3* promoter region (approximately −1000 to −1 bp upstream of the transcription start site), highlighting predicted *cis*-regulatory elements. NAC transcription factors recognize and bind to degenerate core motifs, primarily CACATG, which are often dispersed across target gene promoters. **B**, Localization of predicted NAC-binding *cis*-elements within the *FUS3* gene body (including introns and exons). **C**, Domain architecture of NAC38 and NAC58, illustrating their conserved NAC DNA-binding domains (annotated using Expasy Prosite; https://prosite.expasy.org). **D**, Multiple sequence alignment of the full-length amino acid sequences of NAC38 and NAC58, emphasizing conservation within the NAC domain.

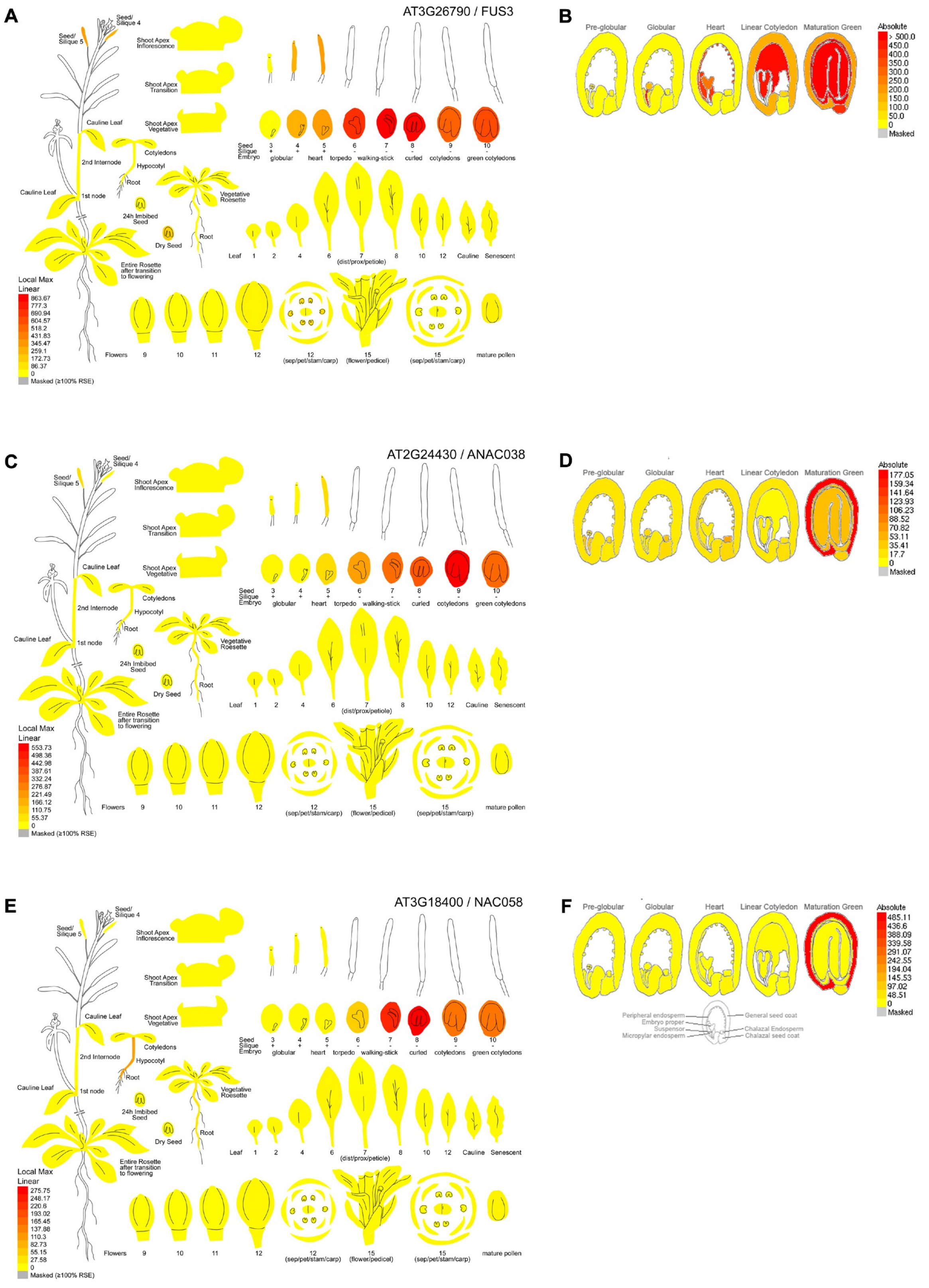


**Supplementary Figure 2. Expression profiles of *FUS3*, *NAC38*, and *NAC58* in *Arabidopsis*.**
**A** to **F,** Images were generated from the BAR eFP browser (https://bar.utoronto.ca; Winter et al., 2007). **A, C, E,** Absolute expression levels of *FUS3*, *NAC38*, and *NAC58* across *Arabidopsis thaliana* developmental stages, based on microarray data from Schmidt et al. (2005). **B, D, F,** Absolute expression levels and heatmaps analysis of *FUS3*, *NAC38*, and *NAC58* in laser-microdissected seed tissues, derived from microarray datasets reported by Le et al. (2010) and Belmonte et al. (2013). NAC38 exhibited the lowest overall expression among the three genes across all analyzed tissues and stages.

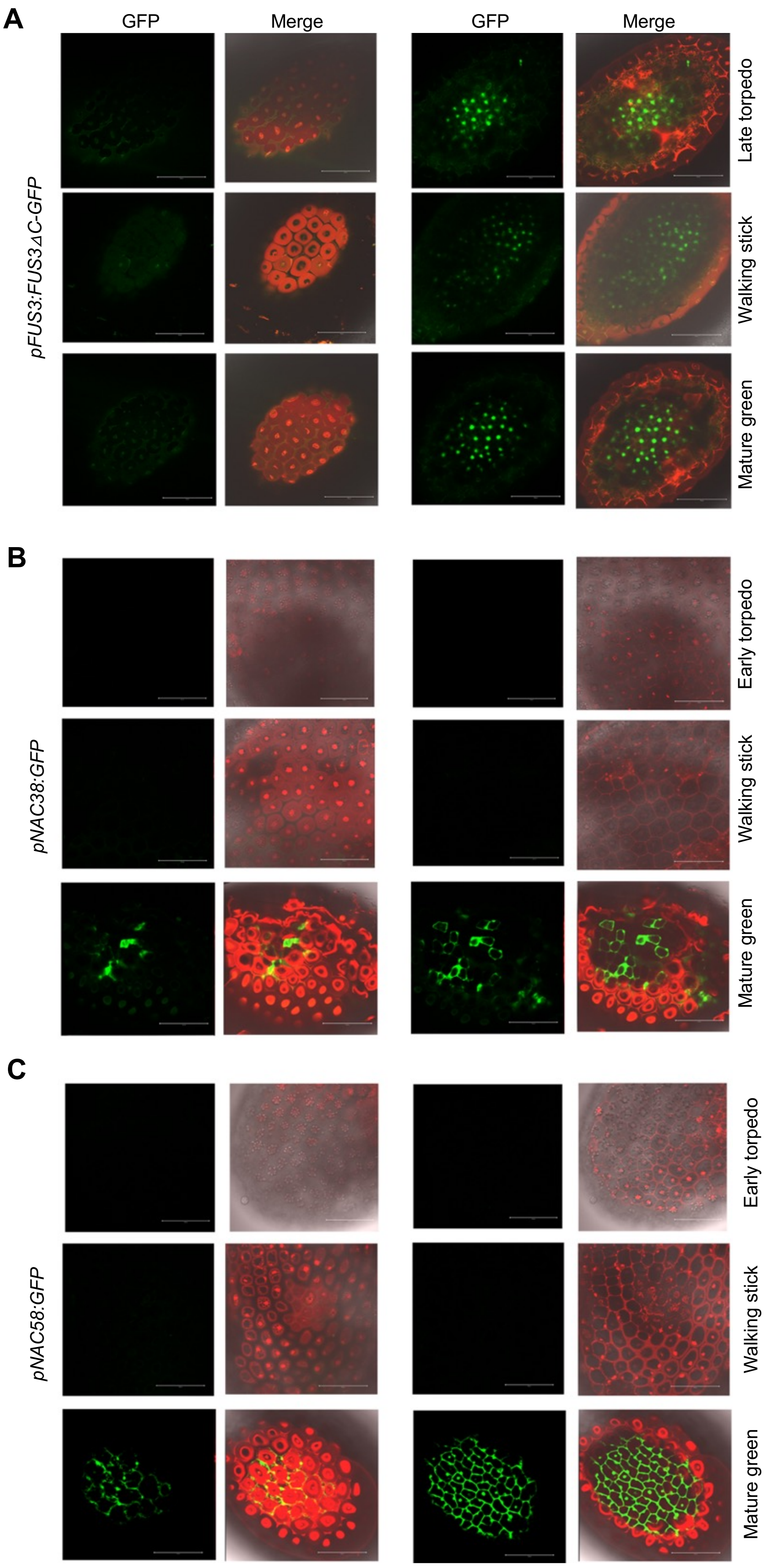

A
GFP
Merge
GFP
Merge
pFUS3:FUS3ΔC-GFP
Late torpedo
Walking stick
Mature green
B
pNAC38:GFP
Early torpedo
Walking stick
Mature green
C
pNAC58:GFP
Early torpedo
Walking stick
Mature green

**D**

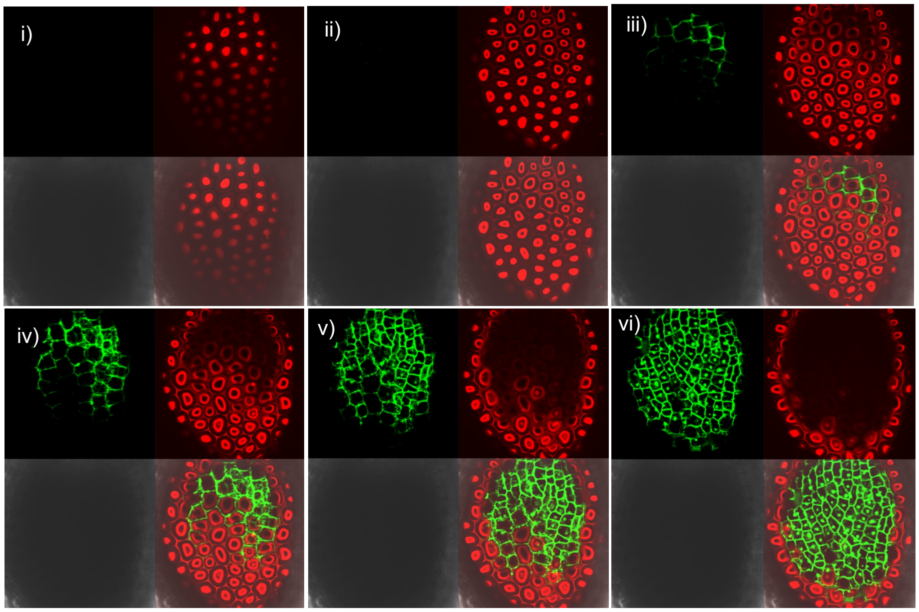


**Supplementary Figure 3. Expression patterns of *FUS3*, *NAC38* and *NAC58* in developing seeds**.

**A** to **C**, Confocal images showing FUS3p:FUS3△C-GFP **(A),** NAC38p:GFP **(B)** and NAC58p:GFP **(C)** fluorescence in the outer integuments (oi) of developing seeds. oi2, left; oi1, right. Scale bars, 100 µm. **D,** Confocal images showing Z-stack series (i to vi) of *NAC58p:GFP* seeds at the mature green embryo stage, from the surface of oi2 inwards towards the oi1, showing GFP in the oi2 (iii, iv) and oi1 (v, vi). Green, red (autofluorescence), merge, and DIC channels are shown clock-wise for each panel in D. GFP was excited at 488 nm and emission was detected at 515-535 nm.

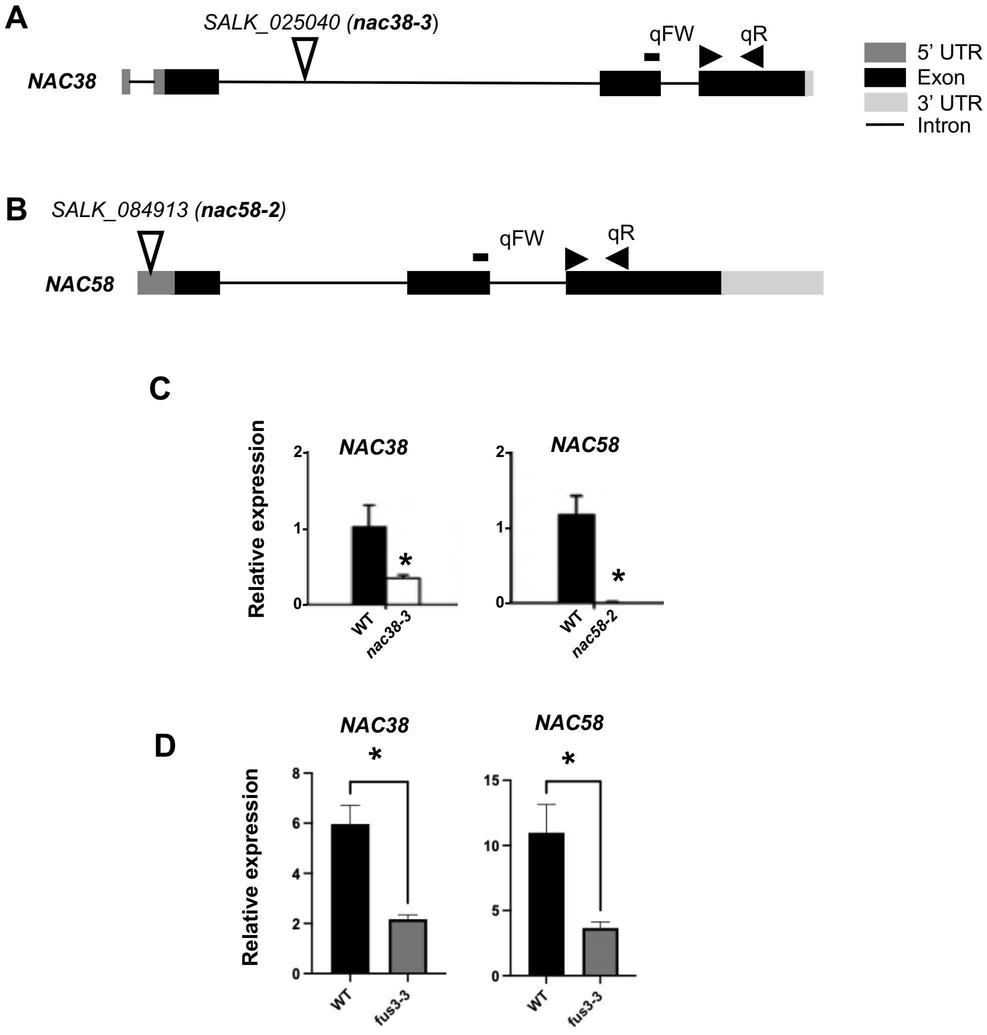


**Supplementary Figure 4. Isolation and molecular characterization of *nac38-3* and *nac58-2* mutants.**

**A, B,** Diagrams showing 5' UTR (dark grey), 3' UTR (light grey), exons (black), introns (black line), and T-DNA insertion sites (white arrows) in *nac38-3* and *nac58-2* mutants. RT-qPCR forward and reverse primer annealing sites are shown by black arrows. **C**, Relative expression levels of *NAC38* and *NAC58* in green seeds of *nac38-3* and *nac58-2* T-DNA insertion mutants by RT-qPCR. **D,** Relative expression of *NAC38 and NAC58* in WT and *fus3-3* 14 DAF seeds by RT-qPCR. Values in C and D represent averages of three biological replicates ± SEM. T-test ($p < 0.05$).

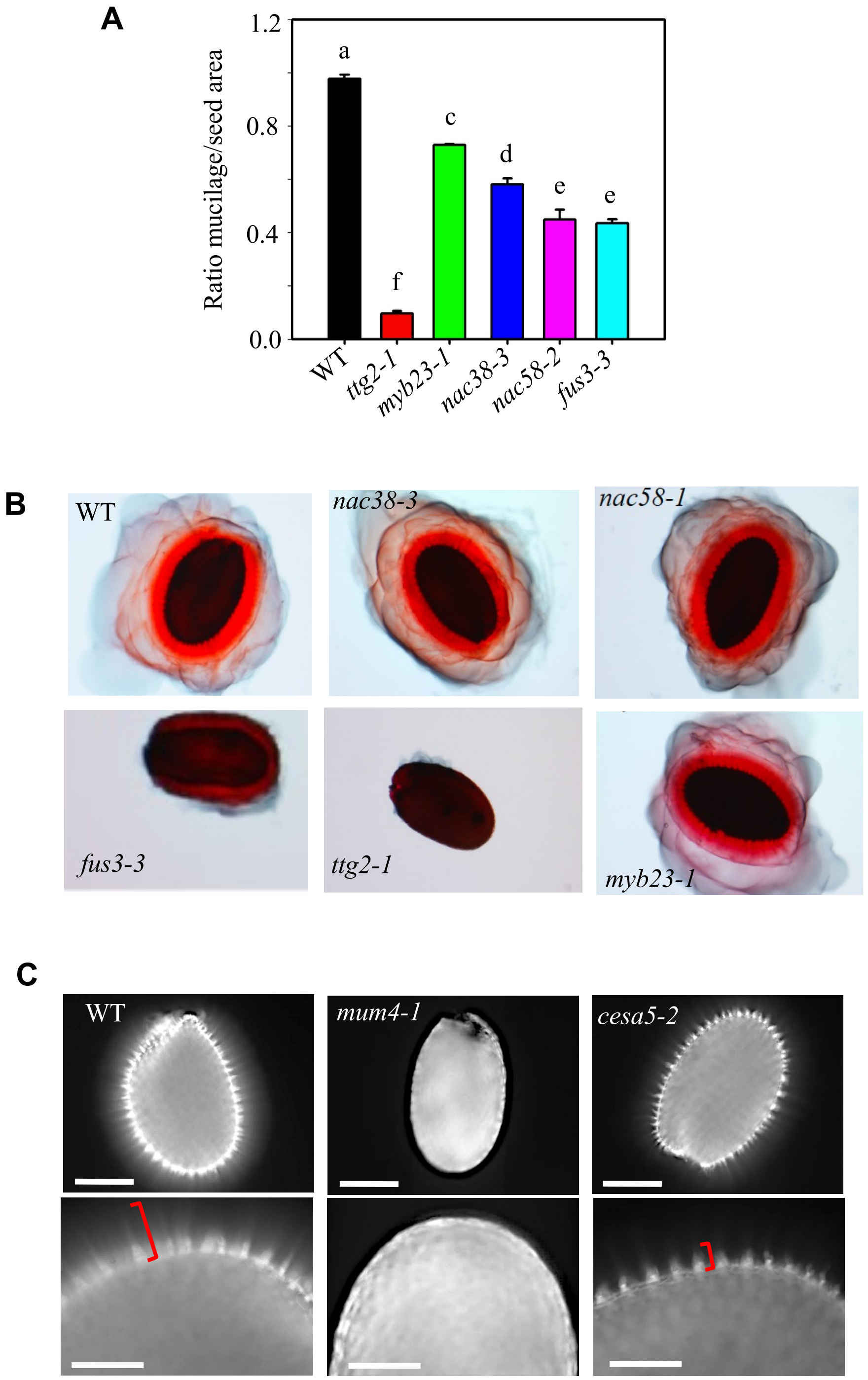


**Supplementary Figure 5. Staining of cellulose and pectin in seed mucilage of WT and mutants.**

**A**, Ratio of adherent mucilage area to seed area of seeds incubated in Ruthenium red with water. Values represent means of three biological replicates ± SEM (n=20-30 seeds per replicate). One-way ANOVA followed by Tukey HST. The *ttg2-1 and myb23-1* were used as controls. *ttg2-1does not* produce mucilage and *myb23-1* has reduced mucilage (Li et al., 2009). **B**, Non-adherent mucilage released from seeds directly stained with Ruthenium red for 30 min **C,** Cellulose staining with Calcofluor white of WT, *mum4-1* and *cesa5-2*. Dry seeds were stained for 1 min and visualized under UV light using Zeiss Epifluorescence microscope. Scale bar = 200 um. Bottom images are close-up of top images. Brackets point to the area of length of cellulosic rays in WT and *cesa5-2*, showing decreased intensity of the diffused stain between the columella and shorter rays of *cesa5-2*. Mucilage is not released in the pectin biosynthesis mutant, *mum4-1 (rhm2)*, as previously shown (Western et al., 2004).

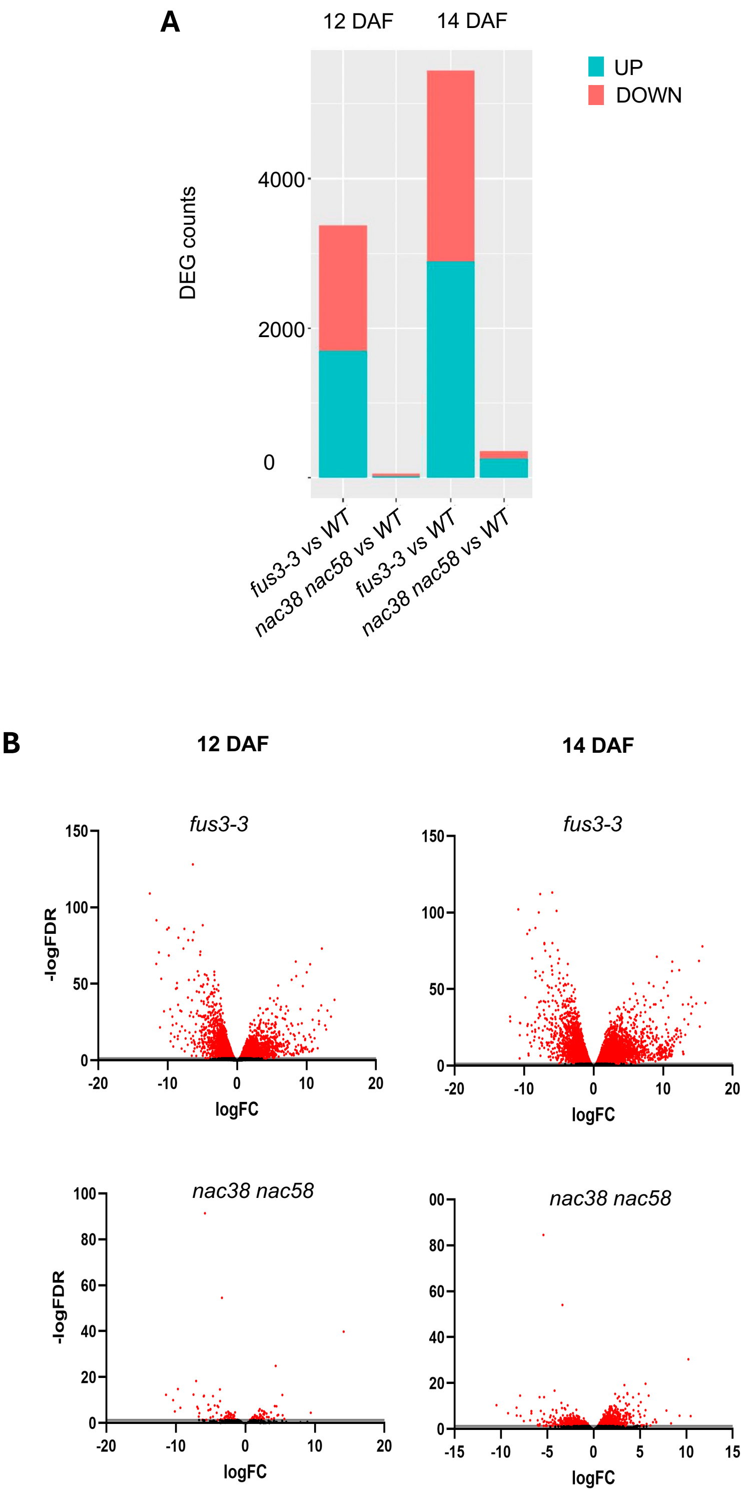


**Supplementary Figure 6. Differentially expressed genes at 12 and 14 DAF in *fus3-3* and *nac38 nac58* by RNA-seq**

**A**) Total upregulated (green) and downregulated (magenta) DEGs in 12 and 14 days after flowering (DAF) seeds. Log2 fold change, FDR 0.05% of DEG are shown. **B)** Volcano plot of differentially expressed genes (DEGs) at 12 DAF and 14DAF.

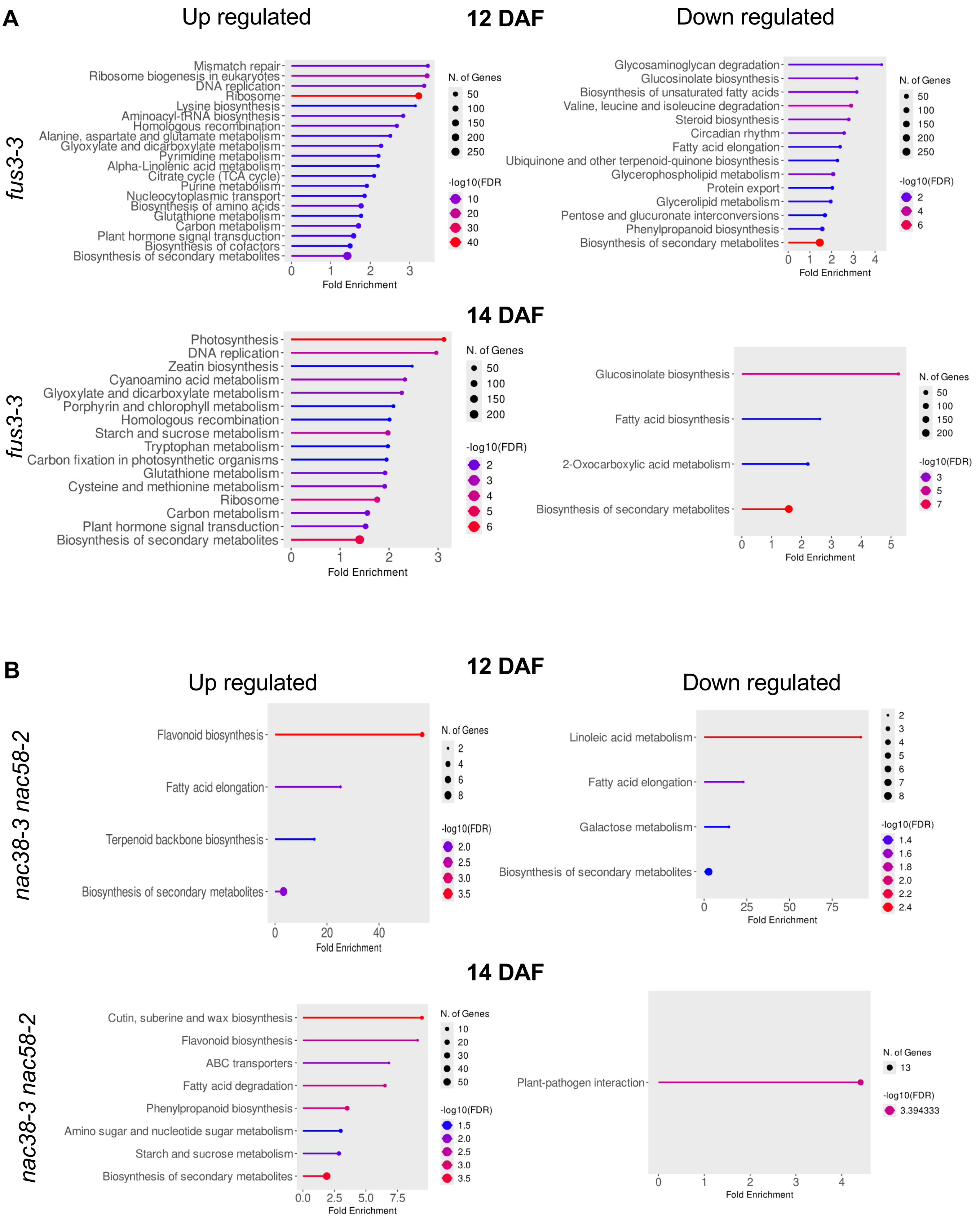


**Supplementary Figure 7. GO biological function enrichment analysis in differentially expressed genes at 12 and 14 DAF in *fus3-3* and *nac38-3 nac58-2***

GO biological function analysis of differentially expressed genes (DEGs) displaying enrichment in up and downregulated genes of *fus3-3* (**A**) and *nac38-3 nac58-2* (**B**) showing the top 10 biological functions at 12 DAF and 14 days after flowering (DAF). Go term analysis performed with ShinyGO (Ge et al. 2020). DEGs includes genes with at least 2-fold change in gene expression and FDR < 0.05.

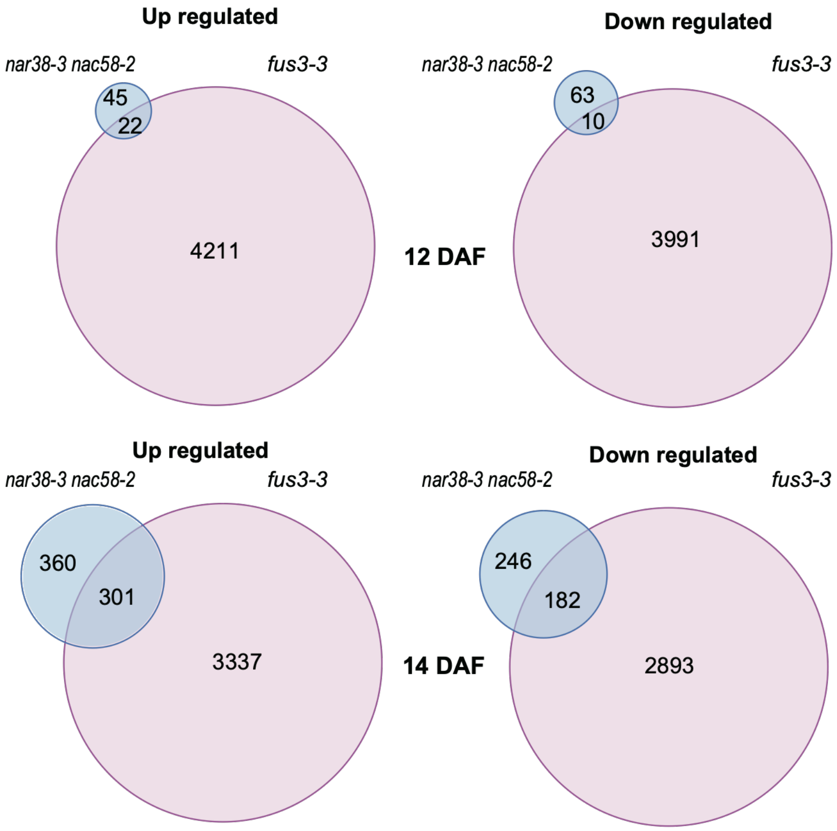


**Supplementary Figure 8. Overlap in differentially expressed genes at 12 and 14 DAF between *fus3-3* and *nac38-3 nac58-2***

Venn diagrams show overlap between *fus3-3* and *nac38-3 nac58-2* DEGs at 12 DAF and 14 days after flowering (DAF). Note that FUS3 is also expressed in the embryo and endosperm (Supplemental Figure S2).

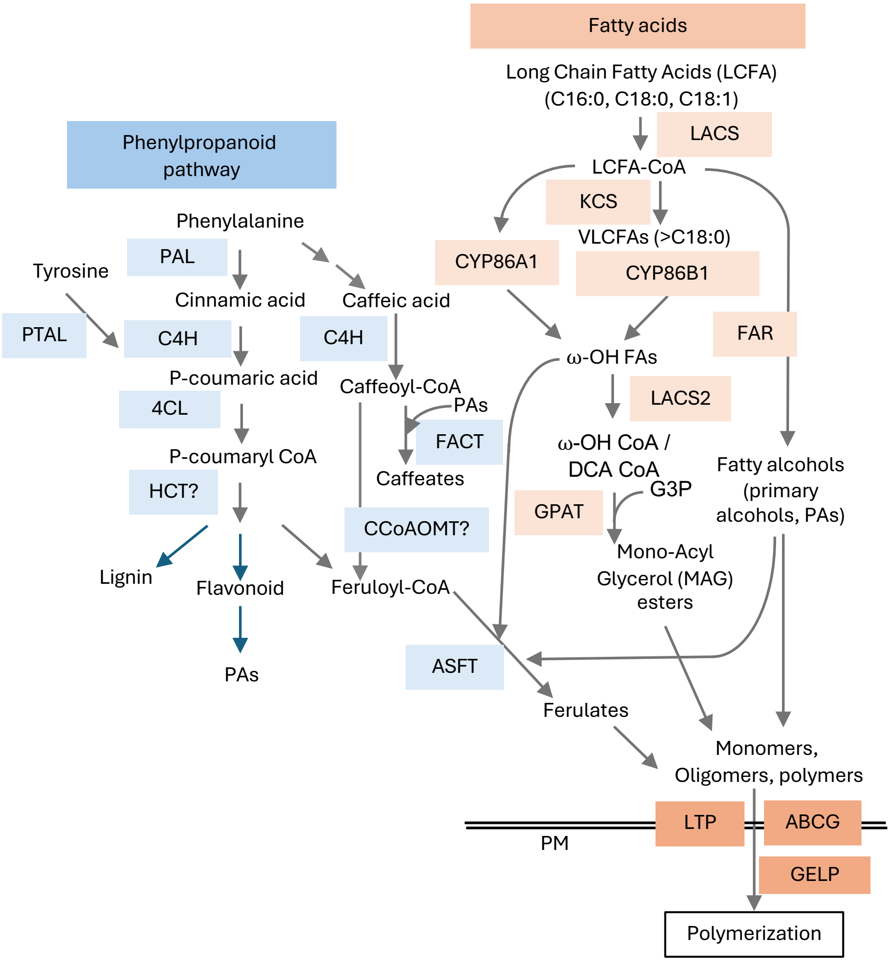


**Supplementary Figure 9. Suberin metabolism pathway**
Drawings showing suberin biosynthesis, transport, and polymerization.

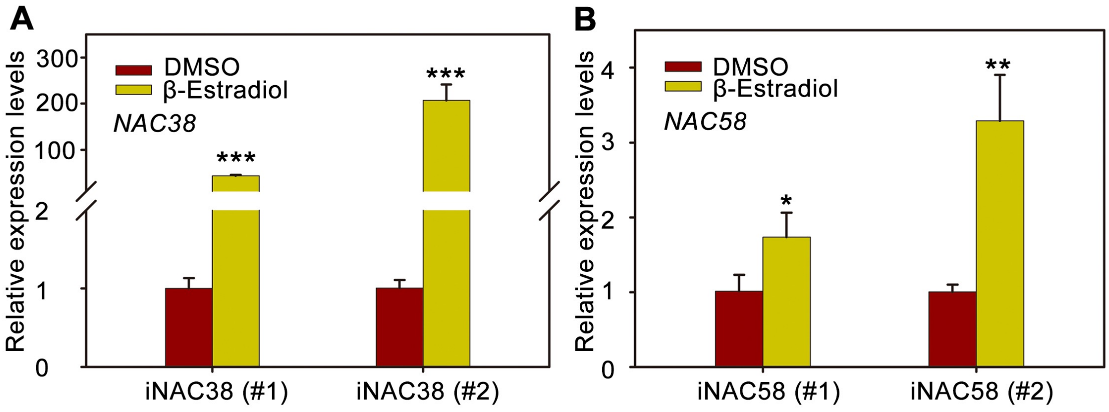


**Supplementary Figure 10 Generation of *iNAC38* and *iNAC58* inducible lines**

Quantitative RT-PCR assessment of *NAC38* (A) and *NAC58* (B) expression levels in β-estradiol-inducible transgenic lines (*iNAC38* and *iNAC58*) following β-estradiol treatment. Data represent means ± SD from three independent biological replicates. ns indicates no statistically significant difference; asterisks denote significance levels determined by Student's *t*-test (* $P < 0.05$; ** $P < 0.01$; *** $P < 0.001$).

A

| Types | NAC38_IP | NAC38_Input | NAC58_IP | NAC58_Input |
|---|---|---|---|---|
| Clean Reads Number | 60,237,872 | 44,909,948 | 59,697,970 | 36,903,994 |
| Clean Q30 Bases Rate(%) | 91.36 | 93.96 | 91.59 | 93.68 |
| Unique Mapped Reads | 39,479,008 | 28,528,453 | 40,232,249 | 21,127,334 |
| Unique Mapped Ratio (%) | 65.54 | 63.52 | 67.39 | 57.25 |

B

Peak height: 0-180

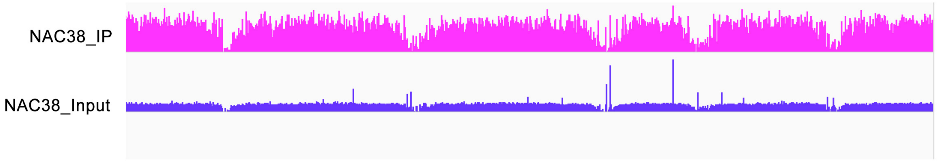


NAC38 ChIP peaks over Chromosomes

C

Peak height: 0-80

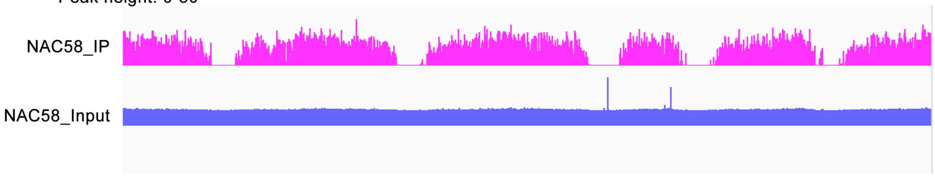


NAC58 ChIP peaks over Chromosomes

D

| Homer *de nove* Motif | *P*-value |
|---|---|
| 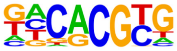 | 1e-122 |
| 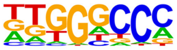 | 1e-56 |
| 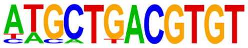 | 1e-48 |
| 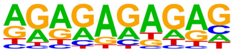 | 1e-46 |
| 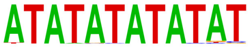 | 1e-39 |

E

| Homer *de nove* Motif | *P*-value |
|---|---|
| 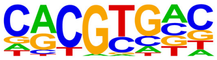 | 1e-100 |
| 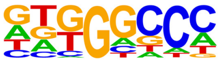 | 1e-82 |
| 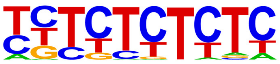 | 1e-46 |
| 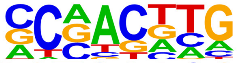 | 1e-36 |
| 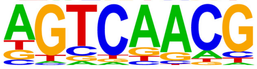 | 1e-36 |

**Supplementary Figure 11. ChIP-seq characterization of NAC38 and NAC58 binding landscapes.**

**A,** Summary statistics of NAC38 and NAC58 ChIP-seq experiments. **B** and **C,** Genome-wide distribution of high-confidence NAC38 and NAC58 binding sites, respectively. **D** and **E,** HOMER-based enrichment analysis of *de nove* binding motifs within NAC38 (**D**) and NAC58 (**E**) peak regions. The top five most significantly enriched motifs (ranked by *p*-value) and are shown.

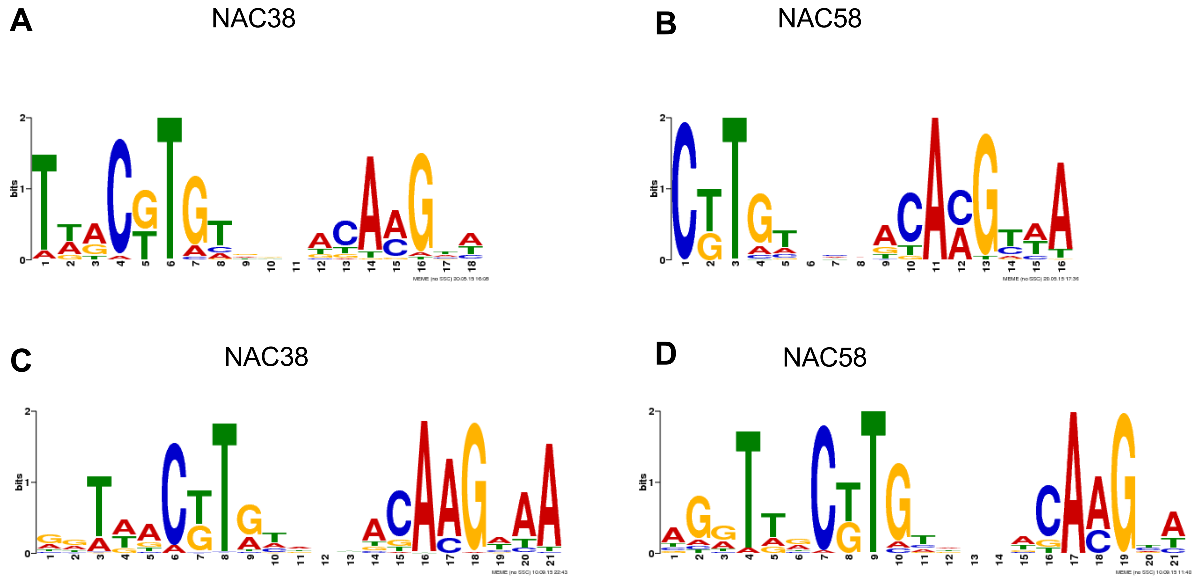


**Supplementary Figure 12. Motifs bound by NAC38 and NAC58 in DAP-seq.**
**A** to **D**, Sequence logo of NAC38 and NAC58 binding motifs were retrieved from DAP-seq (**A,B**) and ampDAP-seq (**C,D**); ampDAP-seq includes binding sites on un-methylated DNA (PCR-amplified genomic DNA, which removes methylcytosines) (O'Malley et al., 2016).

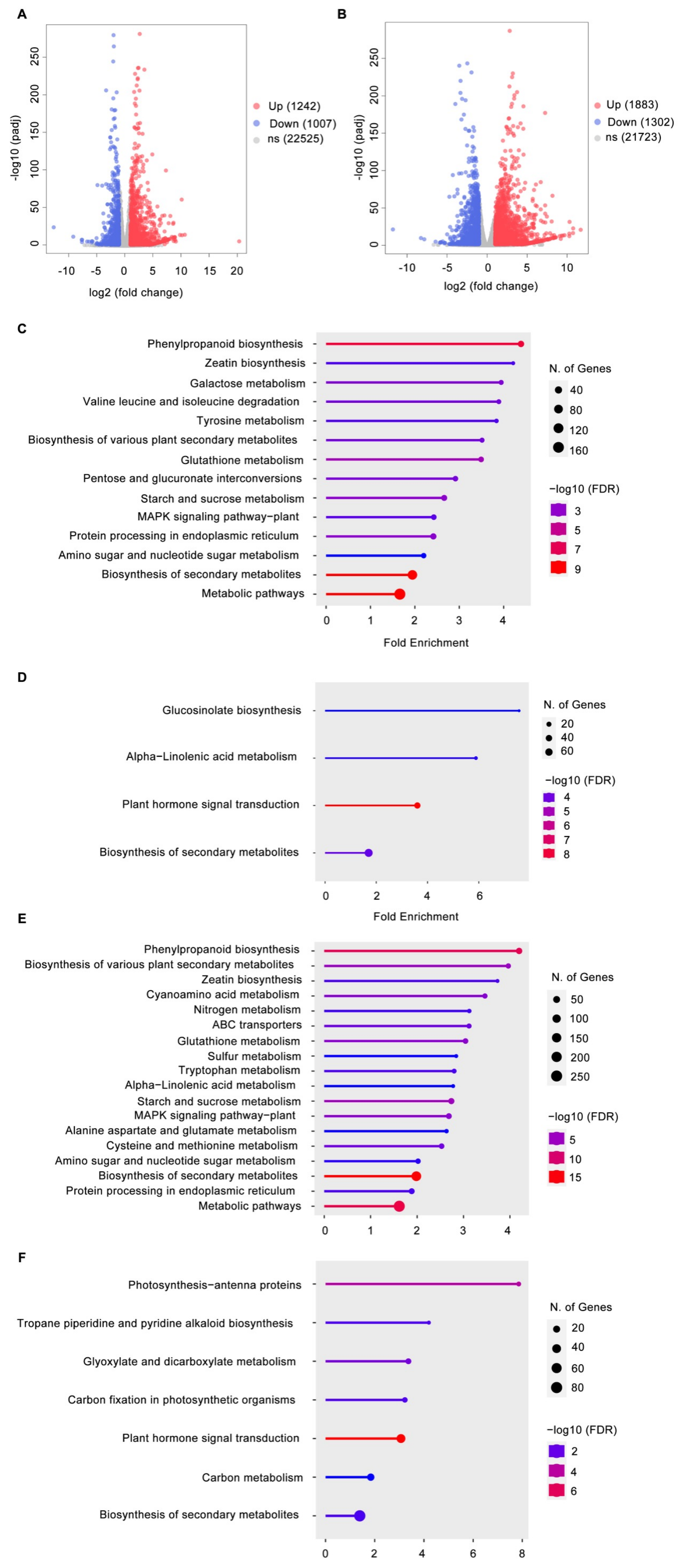


**Supplementary Figure 13. Identification of DEGs in *iNAC38* and *iNAC58* inducible lines.**
**A,** The volcano plots for the DEGs in *iNAC38* inducible lines treated with estradiol compared to DMSO. **B**, The volcano plots for the DEGs in the *iNAC58* treated with estradiol compared to DMSO. **C** and **D**, GO term analysis on biological function displaying enrichment in either upregulated or downregulated genes in *iNAC38* treated with estradiol compared to DMSO, respectively. **E** and **F**, GO term analysis on biological function displaying enrichment in either upregulated genes or downregulated DEGs in *iNAC58* treated with estradiol compared to DMSO, respectively.

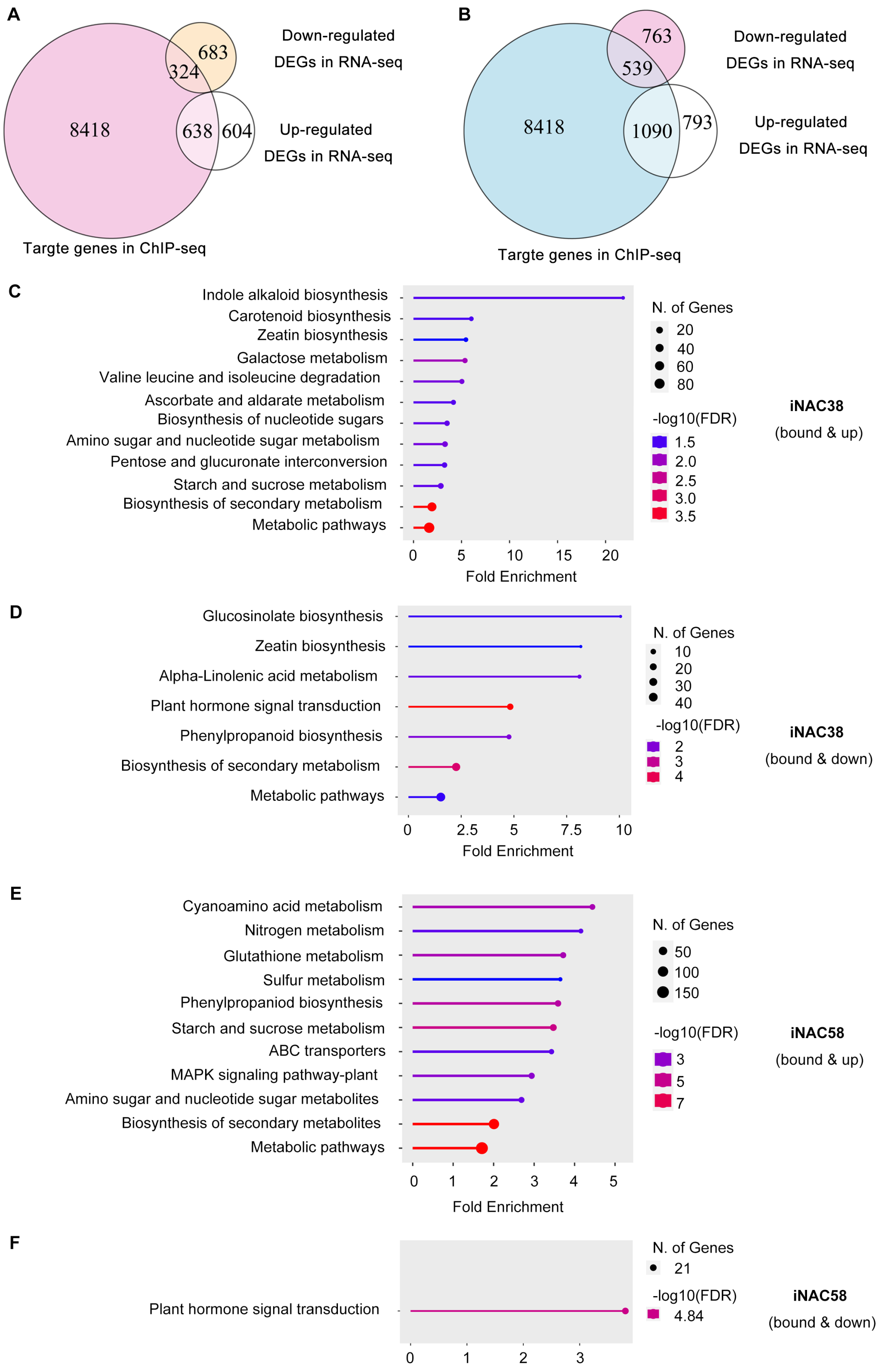


**Supplementary Figure 14. Identification of NAC38 and NAC58 direct target genes (bound and differentially expressed).**

**A** and **B**, Venn diagram showing the number and overlapping genes between the NAC38 or NAC58-binding targets revealed by ChIP-seq and the DEGs identified by RNA-seq in *iNAC38* and *iNAC58* inducible lines, respectively. **C** and **D**, GO term analysis on biological function displaying enrichment in either overlapping upregulated or downregulated genes between ChIP-seq and RNA-seq of *iNAC38*, respectively. **E** and **F**, GO term analysis on biological function displaying enrichment in either overlapping upregulated or downregulated genes between ChIP-seq and RNA-seq of *iNAC58*, respectively. GO term analysis from ShinyGO (Ge et al. 2020).
